# Dynamics of microcompartment formation at the mitosis-to-G1 transition

**DOI:** 10.1101/2024.09.16.611917

**Authors:** Viraat Y. Goel, Nicholas G. Aboreden, James M. Jusuf, Haoyue Zhang, Luisa P. Mori, Leonid A. Mirny, Gerd A. Blobel, Edward J. Banigan, Anders S. Hansen

## Abstract

As cells exit mitosis and enter G1, mitotic chromosomes decompact and transcription is reestablished. Previously, Hi-C studies showed that essentially all interphase 3D genome features including A/B-compartments, TADs, and CTCF loops, are lost during mitosis. However, Hi-C remains insensitive to features such as microcompartments, nested focal interactions between *cis*-regulatory elements (CREs). We therefore applied Region Capture Micro-C to cells from mitosis to G1. Unexpectedly, we observe microcompartments in prometaphase, which further strengthen in ana/telophase before gradually weakening in G1. Loss of loop extrusion through condensin depletion differentially impacts microcompartments and large A/B-compartments, suggesting that they are partially distinct. Using polymer modeling, we show that microcompartment formation is favored by chromatin compaction and disfavored by loop extrusion activity, explaining why ana/telophase likely provides a particularly favorable environment. Our results suggest that CREs exhibit intrinsic homotypic affinity leading to microcompartment formation, which may explain transient transcriptional spiking observed upon mitotic exit.

## INTRODUCTION

3D genome structure and function are variably linked throughout the cell cycle^1^. During mitosis, the nuclear envelope breaks down, chromosomes compact ∼1.5-3-fold, and transcription is largely shut off^1–11^. Condensin II binds chromatin in prophase and extrudes large ∼400-450-kb sized loops, whereas condensin I binds later and extrudes smaller ∼70-90-kb sized loops nested within the larger loops^12,13^. Combined with the loss of CTCF and cohesin from mitotic chromosomes, this largely eliminates all Hi-C-observable interphase 3D genome structural features including TADs, structural CTCF/cohesin loops and functional loops between *cis*-regulatory elements (CREs) such as enhancers and promoters^12,12–16^. A/B-compartments are also lost at this stage^12,15–17^. As cells exit mitosis, interphase 3D genome structures and transcription must therefore be faithfully re-established. Recent work using Hi-C has demonstrated that starting in ana/telophase, A/B-compartments, TADs, and CTCF/cohesin loops form slowly and gradually strengthen to reach full strength by late G1^12,15–20^. Hi-C also detected the dynamics of low connectivity CRE loops, including a small subset of transient ana/telophase specific CRE loops that dissolve again in early G1^1,16,20–22^. However, most CRE loops are poorly resolved by Hi-C^23^ raising the question of how they are dynamically formed at the mitosis-to-G1 transition.

To overcome the detection limits of Hi-C, we recently developed Region Capture Micro-C (RCMC)^24^. RCMC combines Micro-C, which is uniquely sensitive to CRE loops^23,25–27^, with a tiling capture step to focus sequencing reads on regions of interest^24,28^. RCMC achieves >100-fold higher depth than what is possible with genome-wide Hi-C/Micro-C. Using RCMC, we discovered previously undetectable highly nested focal interactions between CREs. We termed these microcompartments because they were largely robust to loss of cohesin-based loop extrusion and appeared to form through an affinity-mediated compartmentalization mechanism akin to block copolymer microphase separation^24^. Thus, microcompartments refer both to a “grid of dots” contact map pattern (nested focal interactions) and a mechanism of interaction (affinity-mediated compartmentalization). Microcompartmental “dots”/loops largely form between CRE anchors, thus appearing as CRE clusters.

Given that all 3D genome structural features are thought to be lost in mitosis, we chose this system to explore the mechanisms and dynamics of microcompartment formation. We applied RCMC to mouse erythroid cells across the mitosis-to-G1 transition. Unexpectedly, we observe microcompartments in mitosis and find that microcompartments transiently peak in strength in ana/telophase before gradually weakening in G1. Integrating 3D polymer modeling, we show how an interplay of affinity, extrusion activity, and chromosome compaction can explain these findings. This provides a mechanistic framework for understanding how loop extrusion, compaction, and affinity-mediated compartmentalization govern 3D genome folding across scales and the cell cycle.

## RESULTS

### RCMC resolves 3D genome folding dynamics from mitosis to G1

To resolve ultra-fine-scale 3D genome folding dynamics following mitosis, we used the experimental system established and validated by Zhang *et al*.^16^. We FACS-purified synchronized mouse G1E-ER4 erythroblasts based on signal from mCherry fused to the cyclin B mitotic degradation domain (mCherry-MD) and on DNA content to achieve ∼98% pure prometaphase (PM), ana/telophase (AT), early-, mid-, and late-G1 (EG1, MG1, LG1) cell populations (**Fig. 1a, Fig. S1,2**). We performed RCMC^24^ to generate deep contact maps at five diverse regions selected for their density of *cis*-regulatory elements (CREs) (**Fig. S1b, 3-6**). Such maps allow us to sharply resolve and follow genomic structures across scales of organization through mitotic exit, including A/B compartments, TADs, and microcompartments which are invisible in sparser datasets (**Fig. 1a, b, Fig. S3-6**).

**Figure 1.**
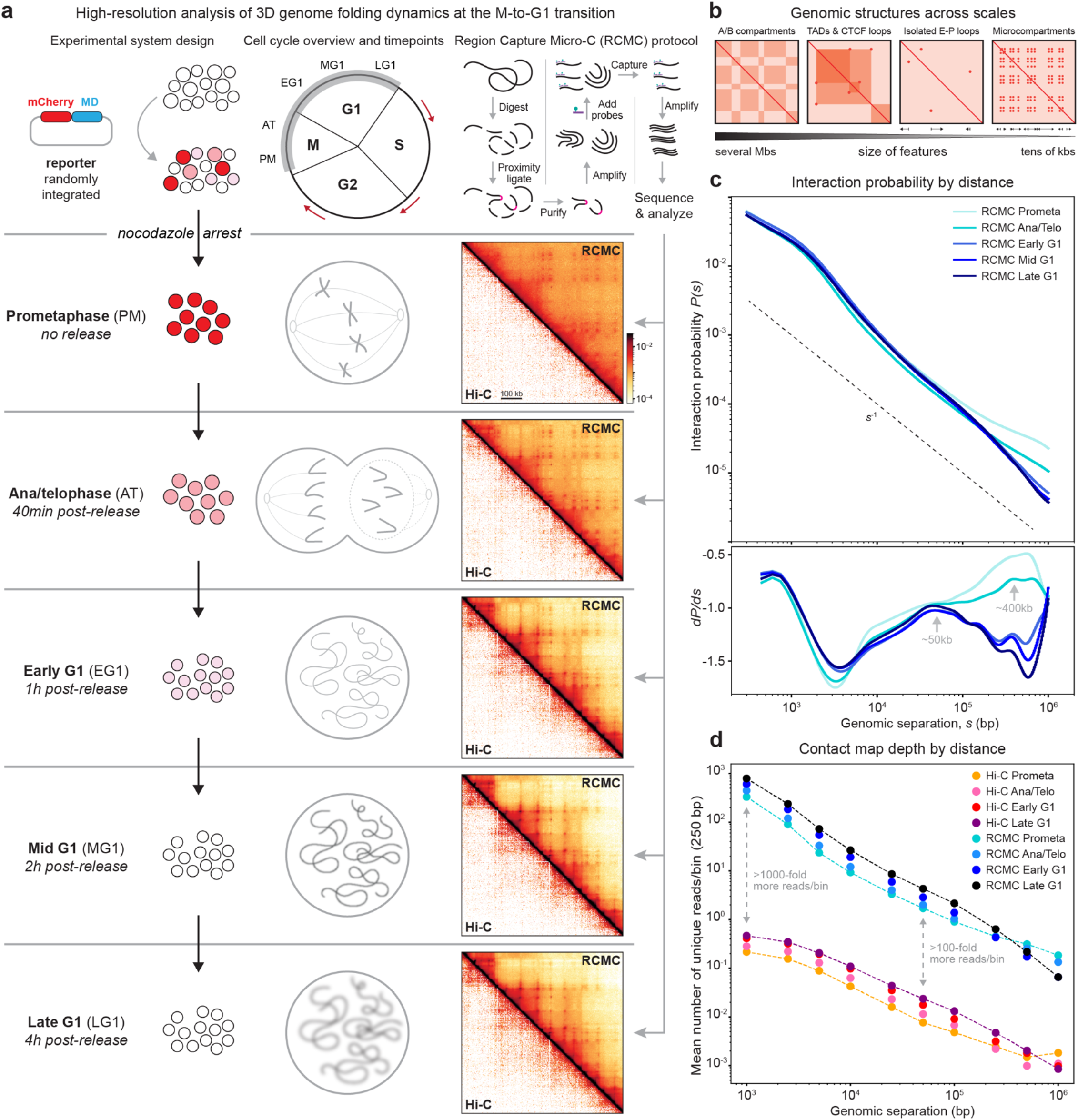
Region Capture Micro-C (RCMC) deeply resolves 3D genomic architecture at the mitosis-to-G1 transition. (**a**) Overview of the experimental system. As previously described^16^, G1E-ER4 cells with an mCherry-tagged mitotic domain reporter are prometaphase-arrested using nocodazole and flow-sorted post-release to capture highly pure cell populations across five mitosis-to-G1 (M-to-G1) timepoints: prometaphase (PM, no release), ana/telophase (AT, 25min post-release), early G1 (EG1, 1h), mid G1 (MG1, 2h), and late G1 (LG1, 4h) (**Fig. S1a**). The Region Capture Micro-C (RCMC) protocol^24^ is applied to each of these cell populations; briefly, chromatin is chemically fixed, digested with micrococcal nuclease (MNase), and biotin labelled before proximity ligation joins spatially proximal fragments. After enrichment for ligated interactions, fragments are library prepped, amplified, and region-captured to create an RCMC library that is sequenced, mapped, and normalized to create contact matrices. (**b**) Schematic representation of how A/B compartments, TADs, CTCF loops, E-P loops, and microcompartments appear in contact maps across scales. (**c**) Interaction probability curves comparing the interaction frequency at different genomic separations (*s*) for the five RCMC datasets. The first derivative of these *P*(*s*) curves is shown at the bottom. (**d**) 3C (Chromosome conformation capture) data density in captured regions for RCMC vs. Hi-C data from Zhang *et al*.^16^. Averaged counts for the number of unique reads across five captured regions are plotted for increasing interaction distances for all datasets at 250 bp bin size.

We obtained the expected interaction scaling with genomic distance, *P*(*s*), for interphase and mitosis^12,15–17^ and observed first derivative peaks of ∼400 kb in mitosis and ∼50 kb in G1 (**Fig. 1c**), which correspond approximately to the average extruded loop sizes^29–32^. Comparing our RCMC maps to prior Hi-C data^16^, we observed the same gradual strengthening of large A/B-compartments and bottom-up formation of TAD and CTCF loops upon mitotic exit thus validating the correspondence between RCMC and Hi-C at coarse resolution (**Fig. S3-6**). Critically, our RCMC maps are between ∼100-fold and ∼1000-fold deeper than the Hi-C data^16^ (**Fig. 1d, Fig. S1c, 2b**) and highly reproducible (**Fig. S1d,e**). This was confirmed by down-sampling the RCMC data by ∼512-1024-fold, which yields comparable data densities (**Fig. S2c**) and contact maps (**Fig. S2d**) to Hi-C^16^. Having validated our RCMC data, we next explored the dynamics of microcompartment formation.

### RCMC reveals nested focal looping interactions between CREs during mitosis

Although similar to Hi-C at coarse resolution, the finer resolutions in our RCMC maps reveal a dramatic restructuring of chromosomes across the cell cycle (**Fig. 2a, Fig. S3-6**). To quantify these dynamics, we began by identifying “dots” in the contact maps, corresponding to focal interactions between two sites (“loops”). We annotated the superset of dots formed across the M-to-G1 transition across the five RCMC regions spanning ∼7 Mb total, yielding 3350 dots between 361 anchors (**Fig. 2a, Fig. S3-6**). Of these 3350 RCMC dots, only 134 were detectable in the Hi-C data (**Fig. S7**). For example, while we identified 888 RCMC dots in the *Klf1* region (**Fig. 2a**), only 20 dots were identified in Hi-C^16^ (**Fig. S7a**). Annotated dots spanned all length scales within captured regions of interest, with a mean length of 368 kb (**Fig. 2b**). Most dots were formed by a subset of high connectivity anchors, with some anchors forming 40-50 clear dots (**Fig. 2c**). To classify dots by their functional identity, we intersected dot anchors with gene promoter annotations (Transcription Start Sites (TSSs)), epigenetic markers of enhancers (H3K27ac and H3K4me1), and structural looping factors (CTCF and the cohesin subunit RAD21), which revealed most anchors to be promoters and enhancers (**Fig. 2d, Fig. S8**). Indeed, we found most dots to be CRE dots/loops (**Fig. 2e**): only ∼1% of dots (34/3350) were “structural loops” lacking CRE overlap, anchored solely by CTCF/cohesin on both sides. Instead, ∼90% of all dots were CRE-anchored on one side and ∼80% CRE-anchored on both sides (P-P, E-P, or E-E). Revisiting the number of dots formed per anchor (**Fig. 2c**) revealed that promoters and enhancers comprise nearly all the high connectivity anchors whereas CTCF/RAD21 anchors form far fewer dots (**Fig. 2f**). The high connectivity of CRE anchors contrasting with CTCF-anchored dots is consistent with a different interaction mechanism for CREs, such as affinity between similar chromatin states and/or transcription factors. Thus, CRE anchors form numerous dots leading to microcompartment formation (“grid of dots” pattern, **Fig. 1b**) unlike CTCF/cohesin anchors which form few dots.

**Figure 2.**
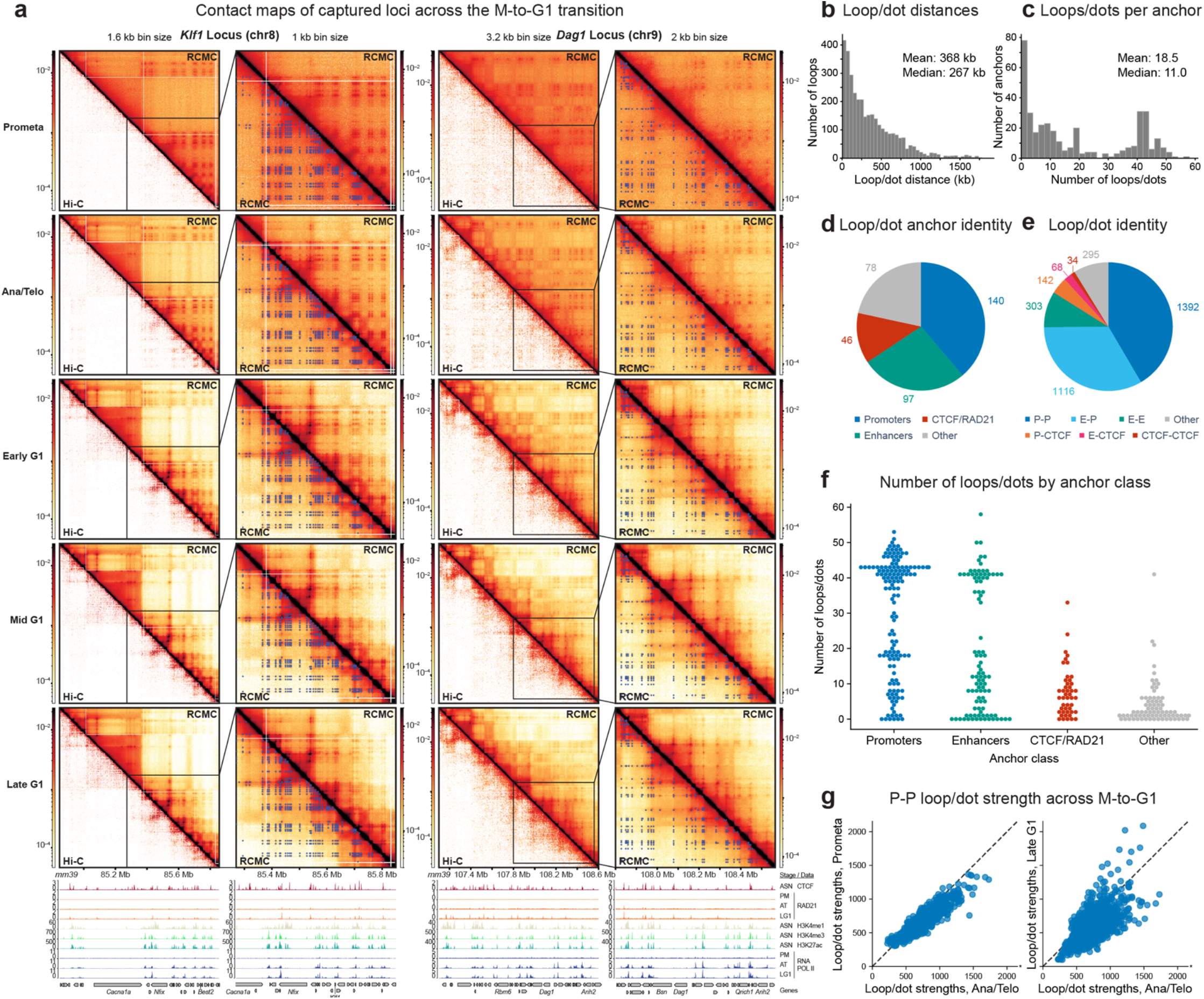
RCMC finely resolves dynamically changing focal looping interactions. (**a**) Contact map visualization of RCMC data at the *Klf1* (bin size: 1.6 kb (left), 1 kb (zoom-in)) and *Dag1* (bin size: 3.2 kb (left), 2 kb (zoom-in)) loci across the M-to-G1 transition, with Hi-C data^16^ (left) and the superset of loops (right) shown below the diagonal. Genomic annotations and ChIP data (stage-specific and asynchronous) are shown at the bottom. (**b-c**) Histograms of (**b**) loop interaction distances and (**c**) the number of interactions formed by each annotated anchor. (**d**) Venn diagram of annotated loop anchors by their genomic identity, determined by chromatin features within 1 kb of anchor sites. Promoters were identified as annotated transcription start sites ±2 kb, enhancers as non-promoter regions with overlapping H3K4me1 and H3K27ac ChIP-seq peaks, and CTCF/RAD21 as non-promoter and non-enhancer sites with overlapping CTCF and RAD21 ChIP-seq peaks. Anchors with multiple overlapping genomic features were hierarchically classified into a single classification, with promoters taking precedence, then enhancers, and finally CTCF/RAD21. Anchors designated as “other” do not overlap promoters, enhancers, nor CTCF/RAD21. (**e**) Venn diagram of annotated loops by the genomic identity assigned in (**d**), with P designating promoters, E designating enhancers, and CTCF designating CTCF/RAD21. (**f**) Swarm plot of the number of interactions formed by each annotated anchor, separated by the genomic categories shown in (**d**). (**g**) Plots of individual P-P (promoter-promoter) interaction strengths in the prometaphase (left) and late G1 (right) conditions, plotted against the strengths in the ana/telophase condition (x-axes). Strengths are calculated as the integrated observed loop signal divided by the expected background signal from local *P(s)* curves (“observed over expected”). This panel shows “exclusive” P-P loops; 186 P-P loops that overlapped with CTCF at one or both anchors were removed; all subsequent loop pileups and quantifications of strength by loop identity similarly omit loops meeting both CRE and CTCF/RAD21 loop types. Axes are truncated for ease of visualization and omit one data point in each plot; in the left (PM vs. AT) plot, this point lies at (2558, 1674), while in the right (LG1 vs. AT) plot, this point lies at (2558, 2746).

Notably, we visually observed striking dynamics of microcompartments (**Fig. 2a, Fig. S3-6**). Microcompartments were already visible in prometaphase, before increasing in strength relative to background in ana/telophase, and then gradually weakening upon G1 entry with many microcompartmental dots being erased by late G1 (**Fig. 1b, 2a, Fig. S3-6**) Quantitative analysis confirmed this observation. The CRE dots that make up microcompartments (P-P, E-P, E-E) peak in strength in ana/telophase (**Fig. 2g, Fig. S9a**). To better characterize the unexpected transience of microcompartments, we next explored the strengths of different loop types across the mitotic-to-G1 transition.

### Microcompartments transiently strengthen, then weaken, across the M-to-G1 transition

To further explore the dynamics of microcompartmentalization, we began by visualizing representative examples of microcompartmental CRE dots (**Fig. 3a, i-iv**) and structural CTCF dots (**Fig. 3a, v**) across the M-to-G1 transition. As above (**Fig. 2a**), the nested CRE dots that comprise microcompartments peak in strength in ana/telophase, in part due to loss of background interaction from prometaphase to ana/telophase, before gradually weakening during G1 (**Fig. 3a, i-iv**). Dot pileups (**Fig. 3b, Fig. S9b**) and strength quantifications (**Fig. 3c, Fig. S9a**) for each functional categorization revealed that CRE dots weaken relative to their background after peaking in ana/telophase, whereas CTCF-anchored dots are relatively weak in prometaphase but monotonically strengthen to be stronger than CRE dots by G1. This observation matches characterizations of CTCF dots from Zhang *et al*.^16^.

**Figure 3.**
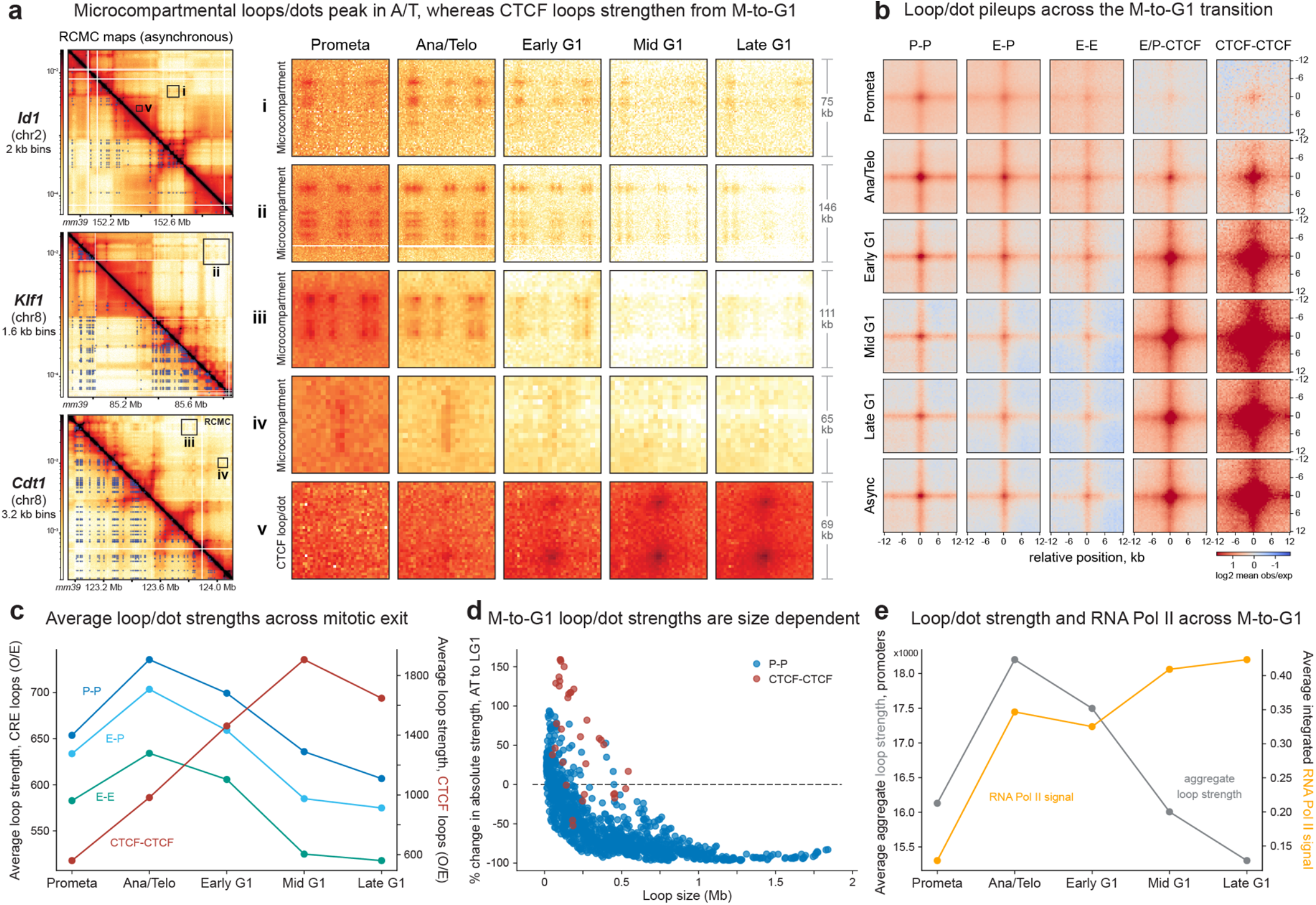
The strength of CRE loops/dots and microcompartments peaks in ana/telophase and weakens as cells enter G1 phase. (**a**) Asynchronous RCMC contact maps (left) at the *Id1, Klf1*, and *Cdt1* regions with manually annotated interactions shown below the diagonal and zoom-in boxes shown in greater detail across the M-to-G1 transition on the right using consistent color map scaling. Zoom-ins show examples of microcompartmental CRE loops in i-iv and CTCF/RAD21 loops in v. (**b**) Aggregate peak analysis (APA) plots of loops, separated to show P-P, E-P, E-E, E/P-CTCF, and CTCF-CTCF loops across the M-to-G1 transition and for the asynchronous condition. Plots show a 24 kb window centered on the loop at 500 bp resolution, and the loops plotted here and in all subsequent panels follow the “exclusive” definition of loop identity as in 2g (CRE sites do not overlap with CTCF). (**c**) Average loop strengths across mitotic exit, with CRE loop strengths on the left axis and CTCF/RAD21-anchored loop strengths on the right axis. Loop strengths were calculated as “observed over expected” signal, as in 2g. (**d**) Change in loop strength across mitotic exit as a function of loop size. The percentage change in absolute loop strength for each P-P (blue) and CTCF-CTCF (red) loop from ana/telophase to late G1 is plotted on the y-axis, while the loop size (or interaction distance) is plotted on the x-axis. Absolute loop strengths are calculated as observed signal without normalization for the expected signal. (**e**) Promoter loop strengths (gray) and RNA Pol II signal (yellow) across mitotic exit, with loop strengths on the left axis and Pol II signal on the right. Loop strengths were calculated as the sum of all observed over expected loop strengths at each promoter and averaged across all promoters. RNA Pol II signal was calculated as the aggregate signal within 1 kb of each promoter-classified loop anchor, averaged across all promoters for each M-to-G1 stage.

Above we have quantified dot strength as signal divided by background (observed/expected, used in **Fig. 3b-c**), but it can also be quantified as signal minus background (observed – expected, used in **Fig. 3d**). As orthogonal validation, we therefore also quantified dot strength by subtracting the background, which confirmed our observation that most CRE dots peak in ana/telophase, unlike CTCF dots (**Fig. 3d**). Notably, these dynamics were highly distance dependent (**Fig. 3d**). While longer-range CRE dots generally strongly weakened after ana/telophase, some short-range CRE dots strengthen from ana/telophase to late G1 (**Fig. 3d**).

We next investigated the relationship between microcompartments and transcription, by comparing CRE dot strength against RNA Pol II ChIP signal at promoters (**Fig. 3e**). We find that the strength of promoter dots peaks in ana/telophase, while RNA Pol II occupancy spikes in ana/telophase before slightly decreasing and then further strengthening in G1. Notably, prior studies uncovered a hyperactive transcriptional state, where around half of all genes, including otherwise silent genes, transiently spike in activity near ana/telophase and early G1^1,33–35^. Our finding that enhancers and promoters, with seemingly low selectivity, form microcompartments that peak in strength in ana/telophase may thus be consistent with this transient spike in transcription during mitotic exit.

### Condensin depletion sharpens A/B compartments but not microcompartments

Previous work has shown that A/B-compartments, formed by large continuous blocks of epigenetically distinct chromatin (∼100s-1000s of kb), strengthen after loss of cohesin-mediated loop extrusion in interphase^36,37^. Recently, to generate a loop-extrusion-free chromatin environment while minimizing the confounding effects of transcription and most transcription factors, we depleted SMC2, a common subunit of condensin I and II, in prometaphase^20^. This led to a very strong gain in A/B-compartmentalization and low connectivity CRE dots in mitosis^20^. These observations prompted us to explore how microcompartments self-organize without condensins.

We applied RCMC to the same experimental system^20^ (**Fig. 4a**). We performed RCMC on prometaphase mitotic chromosomes across five SMC2 depletion timepoints, with deeply resolved contact maps for the 0h, 1h, and 4h depletion timepoints (**Fig. 4b**) and sparser datasets for the 0.5h and 8h timepoints (**Fig. S10-14**). We observe visually striking strengthening of contrast in the checkerboard pattern characteristic of A/B compartmentalization after condensin depletion (**Fig. 4b**). The strengthening of large-scale A/B compartments matches what we previously observed^20^, thus validating our RCMC maps.

**Figure 4.**
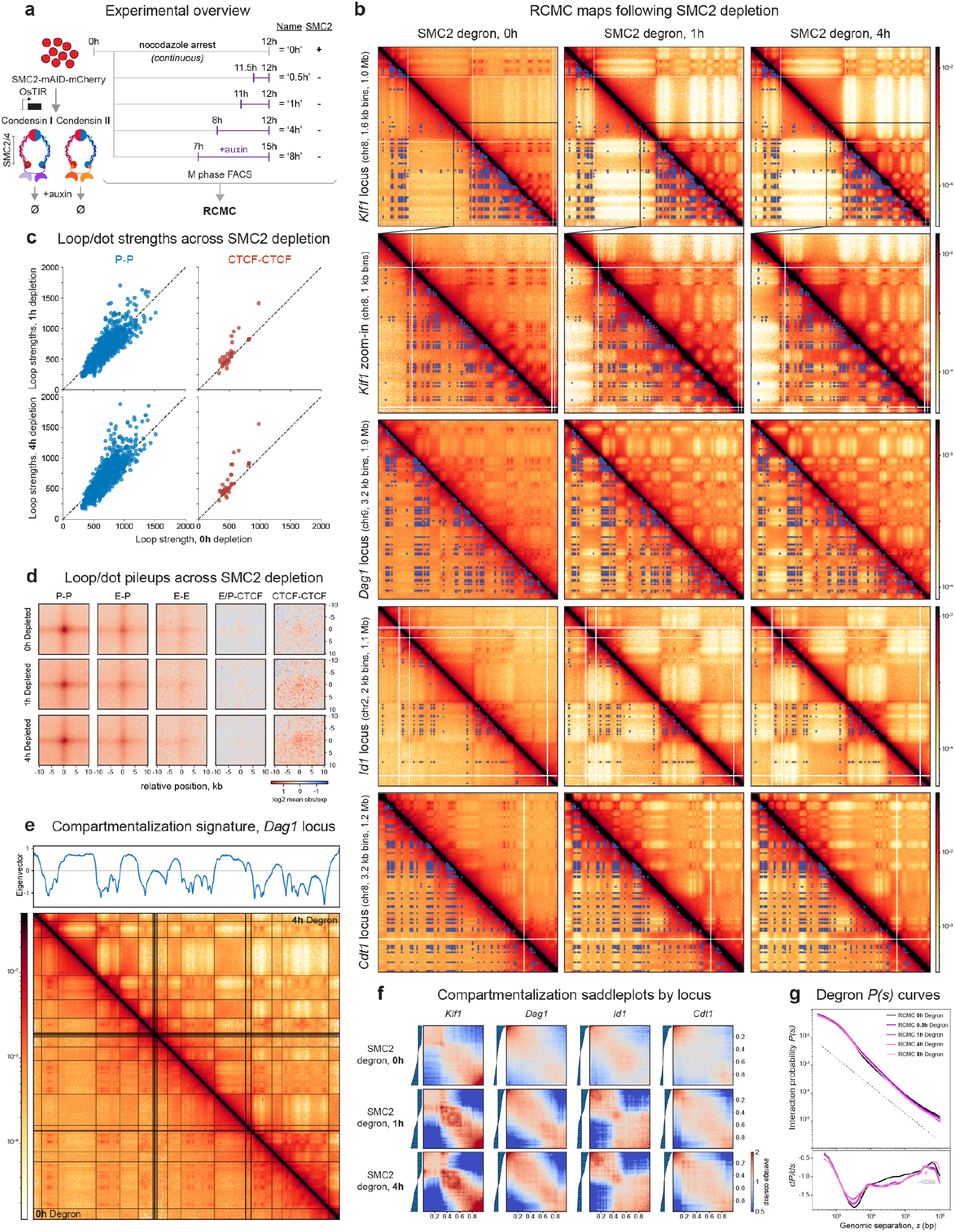
Condensin depletion sharpens A/B compartmentalization while preserving microcompartments. (**a**) Overview of the experimental system. As previously described^20^, G1E-ER4 cells with mCherry-tagged SMC2-mAID are prometaphase-arrested using nocodazole and treated with auxin to induce rapid depletion of SMC2 for 0h, 0.5h, 1h, 4h, and 8h at the end of an 8h (all but the 8h depletion) or 15h (the 8h depletion) nocodazole arrest. SMC2 degradation eliminates both condensin I and II. Cells are then RCMC crosslinked, sorted for M phase purity, and processed into sequencing libraries using the RCMC protocol.(**b**) RCMC contact maps comparing the *Klf1* (plus zoom-in), *Dag1, Id1*, and *Cdt1* loci following 0h, 1h, and 4h of SMC2 degradation. Interaction annotations generated from the M-to-G1 RCMC data are overlaid below the diagonal, and contact intensity scaling is shown on the right. (**c**) Plots of individual loop strengths in the 1h (top) and 4h (bottom) depletion conditions, plotted against the strengths in the control (0h) depletion condition (x-axes), for P-P loops (left) and CTCF/RAD21-CTCF/RAD21 loops (right). Loops were defined by their exclusive identities (no CRE & CTCF overlap) and strengths were calculated as observed over expected signal. (**d**) APA plots of called interactions, separated to show exclusively-defined P-P, E-P, E-E, E/P-CTCF, and CTCF-CTCF loops across SMC2 depletion. Plots show a 20 kb window centered on the loop at 500 bp resolution. (**e**) Compartmentalization signature for the 4h SMC2-depletion condition at the *Dag1* locus. Eigenvector decomposition of interaction frequencies is shown above the contact map, with transition states between positive and negative values noted as black lines overlaid atop the RCMC map, and the 4h depletion condition is shown above the diagonal while the control (0h) depletion is shown below the diagonal. (**f**) Saddleplots of progressive compartmentalization across the 0h, 1h, and 4h depletion conditions at the *Klf1, Dag1, Id1*, and *Cdt1* loci. A track showing the strengths of the two compartments and their transition point is shown to the left of the saddleplots, in which B-compartmental regions (e.g., low eigenvector values) are shown towards the bottom and left of the plots while A-compartmental regions (e.g., high eigenvector values) are shown on the top and right. (**g**) Interaction probability curves comparing the interaction frequency at different genomic separations (*s*) for the five condensin depletion datasets. The first derivative of these *P*(*s*) curves is shown at the bottom.

Next, we explored the effects of condensin depletion on microcompartments and quantified individual dot strengths (**Fig. 4c, Fig. S13c**) and pileups (**Fig. 4d, Fig. S13d**) averaged across all dots. While we did observe strengthening of several CRE dots after condensin depletion (**Fig. 4c**), on average the changes to microcompartment dots upon condensin depletion were minor (**Fig. 4d**). In contrast, analysis of large A/B compartments further confirmed that they sharply increase in strength over time^20^ (**Fig. 4e,f**) without strongly affecting microcompartments. Collectively, the condensin depletion RCMC data point towards mitotic loop extrusion acting more antagonistically towards A/B-compartments formed by larger (∼100s-1000s of kb) blocks than towards microcompartments formed by smaller blocks (∼1-10 kb dot anchors), suggesting that the relative sensitivity of compartments to loop extrusion may be size-dependent.

In summary, we find that large A/B-compartments and microcompartments appear to be at least partially mechanistically separable, as they exhibit temporally distinct formation dynamics upon mitotic exit and distinct sensitivities to loss of condensin in mitosis (**Fig. 4g**). To further explore their mechanistic basis, we turned to experimentally constrained 3D polymer simulations.

### Loop extrusion activity, chromatin affinity, and compaction regulate microcompartments

To investigate the biophysical factors underlying formation, maintenance, and dynamics of microcompartments, we developed a polymer model incorporating major mechanisms of chromatin organization^38^. We modeled loop extrusion by dynamically exchanging SMC complexes (condensin and cohesin) that bind to the chromatin fiber and perform two-sided extrusion before unbinding^31,39,40^ (**Fig. 5a**, top). We also modeled affinity-based homotypic interactions for three types of chromatin (A, B, and C) to capture the formation of large A and B compartments and small microcompartments (denoted as type C, for CRE anchors; **Fig. 5a**, bottom left). We specifically modeled the *Dag1* locus, which we embedded in a larger polymer chromosome confined to a sphere at a chosen volume density (**Fig. 5a**, bottom right).

**Figure 5.**
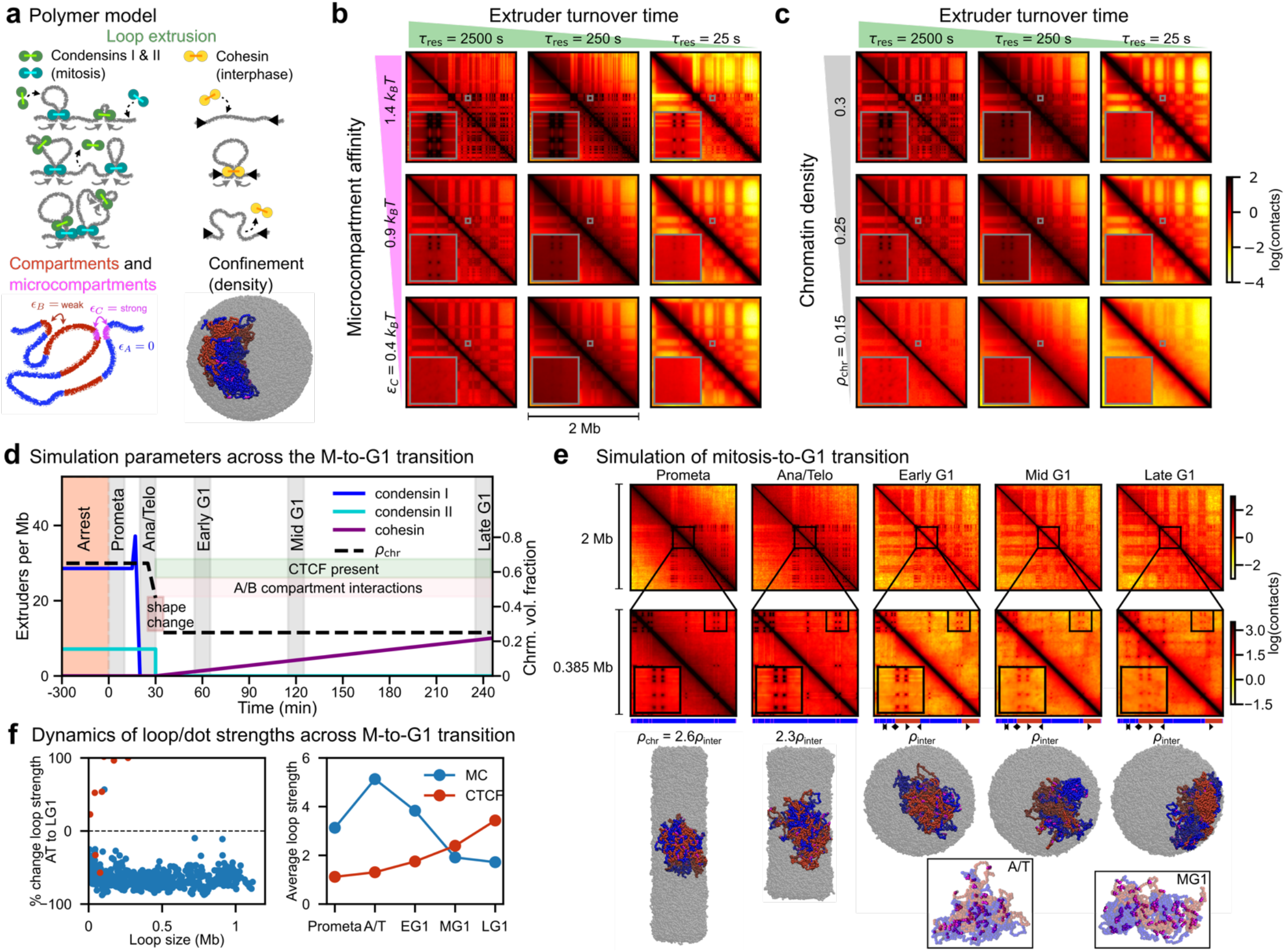
Polymer simulations of chromosomes demonstrate how loop extrusion, interaction energy, and polymer properties may govern microcompartmentalization throughout the mitosis-to-G1 transition. **(a)** Illustrations of key components of the simulation model. Top left: Condensins I and II (green and turquoise, respectively) dynamically bind and unbind to the chromatin fiber (gray) and extrude chromatin polymer loops. Condensin I has a relatively short residence time, τ_res_, which results in the formation of small loops nested within large loops formed by condensin II. Top right: cohesin (yellow) extrudes loops and may stop when it encounters correctly oriented CTCF (black arrowheads). Bottom left: The chromatin fiber is a block copolymer with three types of blocks, which self-interact with affinities given by the interaction energies, *∈*_i_. Bottom right: The *Dag1* region (colored) is simulated as part of a larger polymer chromosome (gray), which is confined to a sphere. **(b)** and **(c)** Contact maps from steady-state simulations of the *Dag1* region for different loop extruder residence times, τ_res_ (decreasing from left to right columns), and **(b)** microcompartmentalization affinities, *∈*_C_ (decreasing from top to bottom rows), or **(c)** different polymer volumetric densities, *ρ*_chr_ (decreasing from top to bottom). Linear density of loop extruders, 1/*d*, is fixed at 1 extruder per 100 kb in these simulations. Small gray boxes denote regions magnified in insets. **(d)** Summary of simulation model of chromosome organization throughout the mitosis-to-G1 transition. Lines show the linear densities of condensins I and II and cohesin, as well as 2.6-fold decrease in polymer density through the mitosis-to-G1 transition. Gray regions indicating time during which data is collected for annotated cell cycle phases. Red region indicates initial equilibration of the simulation, modeling prometaphase arrest. See main text for additional details. **(e)** Contact maps from various times in the mitosis-to-G1 transition simulations with corresponding simulation snapshots (bottom). Middle row displays zoomed-in views of the region indicated in the top row. Insets within this row are the 40 kb x 40 kb region of the contact map indicated by the small black box, showing the dynamics of microcompartmental contacts throughout the transition. Compartment structure and CTCFs are indicated for this region beneath the maps. Images show snapshots of polymer simulation with a single *Dag1* region colored. Boxed images at bottom show snapshots of a 0.385 Mb segment of the *Dag1* region with A and B monomers (blue and red) made transparent to highlight microcompartments (magenta). **(f)** Quantification of percent change in loop/dot strength of simulated microcompartments (MC) from ana/telophase to late G1 as a function of loop size (left) and average microcompartment and CTCF loop/dot strengths (right) throughout the mitotsis-to-G1 transition, as in **Fig. 3(c)-(d)**.

To understand how affinity-based interactions and loop extrusion influence microcompartmentalization, we performed parameter sweeps without extrusion-stalling CTCF sites and computed steady-state contact maps (see **Table 1)**. As microcompartment affinity, *∈*_*C*_, was increased, microcompartments became more visible and more sharply defined (up to ∼8-fold difference from bottom to top row; **Fig. S15a**) because stronger affinity promotes longer-lived interactions (**Fig. 5b**). Furthermore, distinct microcompartments formed in either the presence or absence of a weaker background of larger-scale A/B compartments (**Fig. S15b**). The simulations indicate that microcompartments can be formed by sufficiently strong affinities between small chromatin segments, and that their prominence in contact maps may be tuned largely independently of larger A/B compartments.

Intriguingly, the appearance of microcompartments in the model was also influenced by loop extrusion dynamics. Since cohesin and condensins have different residence times, we tested this by modulating the extruder residence time, τ_res_, at fixed linear density of extruders. We found that faster turnover, i.e. shorter residence time, partially or fully suppressed microcompartmentalization, even for the strongest microcompartment affinities (**Fig. 5b**). Thus, a longer residence time of loop extruders, such as condensin II, results in stronger microcompartments (**Fig. 5b**). This contrasts with previous experimental and computational findings for larger A/B compartments, which can be erased by extrusive cohesin with a long residence time (e.g., due to WAPL loss)^36,41,42^. Instead, for microcompartments, the increase in total extrusion activity (*i*.*e*., extrusion steps per unit time per Mb) induced by faster extrusion turnover can erase or suppress microcompartmentalization (**Fig. S15e**). The notion that extrusion activity is the key to suppressing microcompartmentalization is further supported by simulations with different extruder linear densities and velocities (**Fig. S15a,c-e**). Intuitively, because extruding through a microcompartmental interaction tends to disrupt it, increasing extrusion activity in the model generally weakens microcompartments. Nonetheless, this finding contrasts with the minimal effect of the condensin degradation experiments (**Fig. 4**), therefore suggesting that additional factors are at work during the M-to-G1 transition.

Because chromatin density changes ∼1.5-3-fold through the mitosis-to-G1 transition^43,44^, we simulated systems with different polymer densities (chromosome compaction). We observed that microcompartments were more prominent in systems at higher density (**Fig. 5c**). In denser systems, such as compacted mitotic chromosomes, the configurational entropy of the polymer is decreased due to the decrease in accessible volume. This reduces the entropic penalty of microcompartment formation, thus favoring the formation of microcompartments in more densely compacted chromosomes. Across all simulated densities, increased loop extrusion activity suppressed microcompartments. The effect of density on microcompartment strength is highly non-linear; for a two-fold increase in density, microcompartment strengths increase by ∼30% (**Fig. 5b**), and strengths can be increased ∼6-fold through another twofold density increase (**Fig. S15a**). These simulations indicate that chromatin polymer density can act as a global physical regulator that influences microcompartment formation through both graded and sharp changes.

Together our simulations found that three factors influence the strength of microcompartments. While homotypic affinities between the anchors and higher chromosome density make them stronger, loop extrusion generally weakens microcompartments, with extruders that turn over faster affecting microcompartments to a greater degree.

### Chromatin density and loop extrusion govern microcompartments in simulations of the mitosis-to-G1 transition

An interplay between affinity, extrusion, and density affect microcompartment strength in steady state, but it was unclear how these factors collectively govern microcompartments in a time-varying context, such as the mitosis-to-G1 transition. We implemented the polymer simulation components depicted in **Fig. 5a**, with time-varying, experimentally estimated extrusion and density parameters to model the progression from prometaphase arrest to late G1 (**Fig. 5d**; see Methods and **Table 2**), while holding microcompartmental affinity constant. Timescales were calibrated similarly to simulations in Gabriele *et al*.^45^, using live-cell locus tracking data to properly integrate polymer dynamics and loop extrusion and model the passage of time between cell cycle phases.

The simulation proceeds with (**Fig. 5d**): 1) initialization and equilibration of the chromatin polymer within cylindrical confinement^12^, with microcompartmental affinities and loop extrusion by condensin I and II during prometaphase arrest; 2) prometaphase; 3) condensin I increase at prometaphase, before gradual removal^7,13,46^; 4) ana/telophase, during which the confining cylinder shortens and widens, and polymer density decreases^43,44,47^; 5) condensin II removal^7,15,48,49^, addition of CTCF and A/B compartment affinities^16^, onset of a gradual crossover from cylindrical to spherical confinement at a lower polymer density^50^, and the onset of gradual addition of cohesin^7,16^; 6) early G1; 7) mid G1; and 8) late G1.

Contact maps for simulated chromosomes for each experimental RCMC timepoint in the mitosis-to-G1 transition showed a complex and evolving architecture, as observed in the experiments (**Fig. 5e**). Focal enrichments indicating microcompartments were visible across all timepoints, and they were particularly strong in prometaphase, ana/telophase, and early G1. A/B compartments, TADs, and CTCF-CTCF loops emerged in early G1 and strengthened through G1 as cohesin was loaded and the chromatin polymer re-equilibrated. Notably, simulations reveal that microcompartments are often formed through multi-way interactions; i.e. focal enrichments typically resulted from microphase separation of 5-10 microcompartmental (C-type) anchors.

With the chosen temporal evolution of density and extrusion dynamics, microcompartmental dots peak in strength in ana/telophase, whereas CTCF dots uniformly increased (**Fig. 5f**), as in the RCMC experiments (**Fig. 3c-d**). Our simulations indicate that microphase separation can generate microcompartments. We note that block co-polymer microphase separation is a polymer-based mechanism, that is distinct from protein-based liquid-liquid phase separation^24,36,51–53^. Furthermore, microcompartments can dynamically change via biophysical mechanisms that act and change during the mitosis-to-G1 transition, alongside constant homotypic affinities of CREs.

Simulations suggest that trends in the observed strengths of microcompartments largely, but not exclusively, emerge due to the difference in chromatin densities between mitosis and G1. In simulations in which chromosome density is held constant, microcompartments are stronger in G1 (**Fig. S16**). As observed in experiments (**Fig. 4**), loop extrusion is not necessary to form microcompartments (**Fig. S17a-c**). However, extruders can diminish microcompartments, as observed with shorter residence times (faster turnover) or more loop extruders (**Fig. S17d**). Furthermore, the timing of condensin I removal, is responsible for the strengthening of microcompartments in ana/telophase relative to prometaphase. In simulations in which condensin I is removed during ana/telophase, microcompartments instead peak in strength during early G1 (**Fig. S18a**); this can be remedied by reducing condensin I turnover in ana/telophase (**Ext. Data 18b**). Otherwise, there is little or no dependence on other model assumptions including changes in the shape of the confinement and A/B compartment interactions (**Fig. S19**). Overall, the model generally reproduces experimental contact maps and loop strengths from mitosis to G1.

In summary, our simulation results show that microcompartments are regulated by at least three distinct biophysical factors: homotypic affinity, chromatin density, and loop extrusion activity. Each of these factors, in turn, can be regulated by distinct mechanisms and pathways.

## DISCUSSION

Chromosomes are dramatically reorganized across the mitosis-to-G1 transition. Prior work using Hi-C has shown that essentially all interphase 3D genome structural features – including A/B compartments, TADs, and loops – are lost in mitosis and gradually reformed during G1^1,12,15,15–19,21,54^. Here we apply RCMC^24^ to the mitosis-to-G1 transition^16^ and achieve ∼100-1000-fold higher depth (**Fig. 1c**). Our RCMC maps are consistent with Hi-C at coarse resolution, but unexpectedly reveal a new and previously unobservable layer of 3D genome structure at fine resolution, most notably microcompartments that are present in mitosis. We observe that not only do many CREs come together to form microcompartments in both prometaphase and ana/telophase, but also that most CRE interactions peak in strength in ana/telophase before weakening upon G1 entry (**Fig. 3**).

The presence of microcompartments in mitotic chromosomes provides new insight into the mechanism of microcompartment formation because the formation mechanism must be compatible with the state of the genome in mitosis. Since transcription is largely shut off in mitosis and RNA Pol II absent, their presence in mitosis confirms that microcompartments do not require transcription to form. This is consistent with prior work that finds only modest quantitative changes to CRE loops upon transcription and RNA Pol II perturbations^1,21,22,24,26,55^. Other candidate mediators of microcompartment formation include chromatin state and histone modifications, as well as chromatin and transcriptional regulators. Promoters and, to a lesser extent, enhancers retain chromatin accessibility during mitosis^56^. Furthermore, CREs retain H3K4me1/3 in prometaphase and H3K27ac to some extent^19,57,58^, and H3K27ac likely plays a mitotic bookmarking role^18,57^. Thus, it appears that microcompartments reflect the epigenetic state of mitotic chromosomes, though more work is required to understand whether the relationship is correlative or causal. Moreover, while transcription factors were historically thought to be absent from mitotic chromosomes^59,60^, recent work has found that some factors remain bound to mitotic chromosomes and thus may also serve a mitotic bookmarking function. These include SOX2^61,62^, TBP^63^, BRD4^57^, ESRRB^64^, NR5A2^64^, GATA1^65^, and many others^59,60,66^. Thus, putative mitotic bookmarking proteins are also candidate mediators of microcompartment formation. Finally, we speculate that rather than being fully mediated by a single factor, microcompartments most likely form through a “strength-in-numbers” mechanism involving the combined affinity-mediated interactions of many factors.

Polymer modeling provides further mechanistic insight and shows that microcompartmentalization is largely controlled by three characteristics: homotypic affinity of microcompartment anchors (such as CREs), dynamics of loop extrusion, and chromatin density. While is it unsurprising that stronger affinity leads to stronger microcompartments, our simulations reveal unexpected effects of loop extrusion and density. For loop extrusion, “extrusion activity” 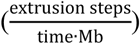 appears to be the key parameter. Each time an extruder such as condensin or cohesin extrudes through a microcompartment anchor, it is likely to disrupt its interactions by bringing other chromatin segments into contact with the microcompartment anchor, regardless of their affinities for each other. Thus, the collective effect of the number of extruders, their residence time, and processivity governs the stability of microcompartments (**Fig. 5b-c, S15a,c-e**). This observation contrasts with large A/B-compartments: for example, increasing extruder residence time via WAPL loss weakens A/B-compartments^36,41,42^. This observation strengthens the notion that microcompartments and larger A/B compartments may be differentially modulated, even though the underlying biophysical mechanisms – affinity plus alterations by loop extrusion – are similar.

Our simulations also reveal that chromosome density/compaction plays an unexpectedly large role: A twofold change in density, which approximately matches the difference between mitotic and interphase chromosomes^43,44^, is sufficient to go from nearly absent to very strongly visible microcompartments (**Fig. 5c**, bottom vs. top rows). Physically, microcompartment formation is favored by enthalpy but disfavored by entropy. Microcompartment formation reduces the configurational entropy of the polymer, but the spatial constraints introduced via compaction reduce this entropic cost, thereby promoting microcompartment formation. Thus, while the presence of microcompartments in mitotic chromosomes was unexpected, their presence is consistent with polymer modeling: microcompartment formation in prometaphase is facilitated by high compaction (**Fig. 5c**), and telophase likely provides a uniquely favorable environment due to the combination of very low extrusion activity^7,13,15^ and high compaction^43,44^ (**Fig. 5d**), thus explaining why microcompartments peak in strength in ana/telophase (**Figs. 3, 5e-f**). This model also predicts that perturbations that affect density (e.g. osmotic shock) may affect microcompartments, and that cell types with higher chromatin density (e.g. smaller nuclei) may form stronger microcompartments. This may also help explain the modest effects of interphase cohesin depletion on E-P interactions^24,67^: because cohesin depletion simultaneously decreases chromosome compaction/density^36,42,45,68^ and decreases extrusion activity, these negative and positive effects on microcompartments may roughly cancel out, thus explaining a relatively modest effect overall^24,67^.

Therefore, chromosome compaction and the lack of loop extrusion by cohesin emerge as leading factors for stronger microcompartments in mitosis. Furthermore, the only consistent models that we found had slow extrusion dynamics after prometaphase (**Figs 5d-f, Fig. S18**). This finding hints at the possibility that mitotic extrusion dynamics after prometaphase may be rather subtle, as high extrusion activity of condensin I would weaken microcompartments. Moreover, the lack of changes in microcompartment strength upon condensin depletion could also suggest that “extrusion activity” by condensins is diminished during later stages of mitosis. Thus, the loops of mitotic chromosomes may be fully extruded by the end of prometaphase with comparably less extrusion activity later.

Our polymer model (**Fig. 5a,d**) reproduces the key features of 3D genome folding during the mitosis-to-G1 transition, including gradual formation of A/B-compartments, TADs, and CTCF loops as well as microcompartments that peak in ana/telophase, but there are several limitations. These include uncertainty about how key parameters change from mitosis to G1, including microcompartment and A/B compartment affinities and extrusion parameters. Additionally, we have not explored the contributions of other mechanisms thought to be involved in A/B compartment formation, such as such as interactions with nuclear bodies (e.g. the lamina, nucleoli, speckles, etc.)^69,70^, chromatin-chromatin crosslinks (e.g. HP1)^71,72^, and active polymer dynamics^73,74^, which might variably facilitate or hinder microcompartmentalization.

The same mechanism of block copolymer microphase separation^52,53,75,76^ might explain compartmentalization across scales: large blocks result in A/B compartments^36^, kb-sized blocks result in microcompartments^24^, and introducing both results in co-existing A/B compartments and microcompartments (**Fig. 5**). This raises the question of whether microcompartments and the active A compartment are formed by the same molecular factors, but at different scales. Several observations from our study suggest that they may be at least partially distinct: 1) microcompartments are strongly visible in mitotic chromosomes, whereas A/B compartments are absent (**Fig. 2-3**); 2) condensin depletion leads to strong A/B-compartmentalization in prometaphase without strongly affecting microcompartments (**Fig. 4**); 3) in simulations, we can largely tune A/B-compartment strength and microcompartment strength independently, without them strongly affecting each other (**Fig. 5, S15a,b** and **S19b-d**); 4) increasing extruder residence time strengthens microcompartments (**Fig. 5b,c**), but weakens A/B compartments^36,41,42^. Thus, although compartmentalization remains poorly understood and much more work is required, our results suggest that microcompartments are at least partially distinct from large A compartments.

Finally, our observation of transiently peaking microcompartments may explain the hyperactive transcriptional state that forms during mitotic exit, during which about half of all genes transiently spike^1,33–35^. While our regions are more gene- and CRE-rich and more work is required to establish generality, our data nevertheless suggests that many CRE interactions (E-P, P-P, E-E) are intrinsically broadly promiscuous and exhibit only moderate selectivity. Indeed, we observe dozens of enhancers and promoters that form >40 distinct dots (**Fig. 2f**), consistent with our prior work in mESCs^24^. Moreover, our polymer model assumes no CRE selectivity and that all CREs have equal affinity for each other, but nonetheless reproduces experimentally observed microcompartmentalization. While our observation of promiscuous CRE interactions leading to the formation of microcompartments does not mean that all or some are causally instructive for transcription, we nevertheless observe CRE interactions slightly precede RNA Pol II promoter binding on average (**Fig. 3e**). Mechanisms that lead to less promiscuous CRE interactions in interphase include chromosome decompaction upon G1 entry, the constraining action of CTCF/cohesin, and perhaps the action of potentially selective CRE looping factors such as YY1 and LDB1^67,80,81^, among other mechanisms. Thus, our data and simulations suggest that CREs are intrinsically broadly interaction-compatible leading to microphase separation-mediated microcompartment formation that peaks in ana/telophase and may explain the broad and transient transcriptional spiking observed during mitotic exit^33^.

## Acknowledgements

We thank Bin Zhang, Joe Paggi, Effie Apostolou, as well as Jacob Kæstel-Hansen, Masahiro Nagano, Clarice Hong, Sarah Nemsick, Miles Huseyin, Michele Gabriele, Sumin Kim, and the Hansen lab for feedback on the manuscript. We also thank Emily Navarrete and the other members of the Mirny Lab for helpful discussions. ASH acknowledges funding support from the NIH (DP2GM140938, R33CA257878, R01EB035127, UM1HG011536, R01CA300848, R03OD038390), an NSF CAREER award (2337728), the Gene Regulation Observatory of the Broad Institute of MIT and Harvard, Pew-Stewart Scholar for Cancer Research, the Mathers Foundation, and an RSC award from the MIT Westaway Fund. This work was supported by the Bridge Project, a partnership between the Koch Institute for Integrative Cancer Research at MIT and the Dana-Farber/Harvard Cancer Center. We thank the MIT Koch Institute’s Robert A. Swanson (1969) Biotechnology Center for technical support, specifically the Integrated Genomics and Bioinformatics Core and MIT BioMicroCenter, and this work was supported in part by the Koch Institute Support (core) Grant P30-CA14051 from the National Cancer Institute. LM and EJB are supported by NIH NIGMS GM114190 and NSF awards 2044895 and 2210558. LM is a Simons Investigator of Simons Foundation. GAB is supported by 5U01DK127405-04. NGA is supported by 1F31DK136200-01A1. We also thank the Walk-Up Sequencing services of the Broad Institute of MIT and Harvard. The data generated in this study can be found at NCBI Gene Expression Omnibus under accession number GSE276657.

## Notes

https://www.ncbi.nlm.nih.gov/geo/query/acc.cgi?acc=GSE276657

### Competing Interest Statement

VYG and ASH have previously filed a patent for Region Capture Micro-C (RCMC). All other authors declare no competing interests.

