## Supplementary Information for "Dynamics of microcompartment formation at the mitosis-to-G1 transition"

[1] Department of Biological Engineering, Massachusetts Institute of Technology; Cambridge, MA 02139, USA; [2] Gene Regulation Observatory, Broad Institute of MIT and Harvard; Cambridge, MA 02139, USA; [3] Koch Institute for Integrative Cancer Research; Cambridge, MA, 02139, USA; [4] Perelman School of Medicine, University of Pennsylvania, Philadelphia, PA, USA; [5] Division of Hematology, The Children's Hospital of Philadelphia, Philadelphia, PA, USA; [6] Institute of Molecular Physiology, Shenzhen Bay Laboratory, Shenzhen, Guangdong, China; [7] Institute for Medical Engineering and Science and Department of Physics, Massachusetts Institute of Technology, Cambridge, 02139 MA, USA

**Contents**

- Supplementary Figures and Figure Legends
- Materials and Methods
- Tables
- Supplementary References

### SUPPLEMENTARY FIGURES AND LEGENDS

**a**

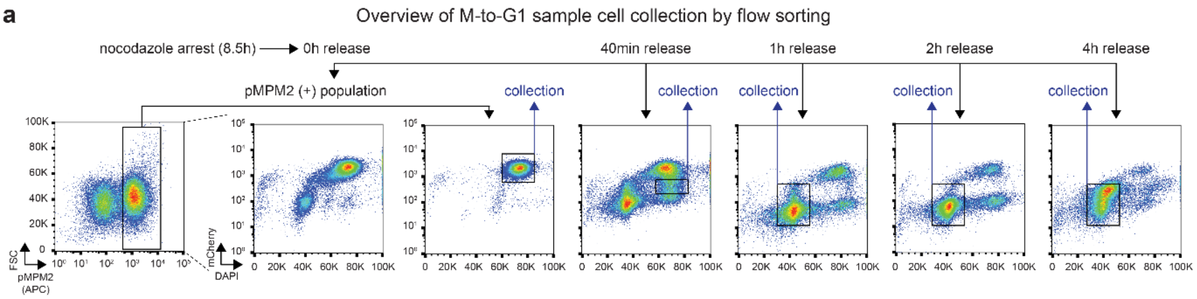


**b** RCMC capture efficiency by locus, asynchronous condition

| | RCMC locus | *Id1* | *Klf1* | *Cdt1* | *Dag1* | *Myc* |
|---|---|---|---|---|---|---|
| | Chromosome | chr2 | chr8 | chr8 | chr9 | chr15 |
| | Coordinates *(mm39)* | 151,920,000–153,000,000 | 84,846,629–85,856,629 | 122,950,000–124,110,000 | 106,675,000–108,600,000 | 61,810,000–63,684,000 |
| | Locus size (Mb) | 1.080 | 1.010 | 1.160 | 1.925 | 1.874 |
| Biological Rep 1 | Mapped reads *(combined capture)* | 546,984,849 | | | | |
| | Mapped reads *(to locus)* | 50,804,677 | 50,915,842 | 67,166,399 | 82,910,072 | 47,419,514 |
| | Capture efficiency | 54.7% | | | | |
| Biological Rep 2 | Mapped reads *(combined capture)* | 448,572,694 | | | | |
| | Mapped reads *(to locus)* | 30,580,165 | 31,836,300 | 40,547,834 | 49,554,868 | 28,707,523 |
| | Capture efficiency | 40.4% | | | | |

**c** Comparison of M-to-G1 data density across RCMC and Hi-C

| Read counts comparison | Locus size (Mb) | Unique reads genome-wide | Unique reads entirely in ROIs |
|---|---|---|---|
| RCMC, asynchronous (async) | N/A | 682,289,559 | 246,293,481 |
| *RCMC, Id1 locus (chr2)* | 1.080 | N/A | 40,123,900 |
| *RCMC, Klf1 locus (chr8)* | 1.010 | N/A | 41,737,270 |
| *RCMC, Cdt1 locus (chr8)* | 1.160 | N/A | 55,698,589 |
| *RCMC, Dag1 locus (chr9)* | 1.925 | N/A | 69,310,808 |
| *RCMC, Myc locus (chr15)* | 1.874 | N/A | 39,422,914 |
| RCMC, prometaphase (PM) | N/A | 586,610,326 | 137,330,275 |
| RCMC, ana/telophase (AT) | N/A | 580,899,254 | 171,485,290 |
| RCMC, early G1 (EG1) | N/A | 684,871,630 | 233,015,417 |
| RCMC, mid G1 (MG1) | N/A | 642,882,309 | 229,120,128 |
| RCMC, late G1 (LG1) | N/A | 754,484,446 | 315,514,810 |
| Hi-C, Zhang et al. (2019), PM | N/A | 379,448,977 | 569,540 |
| Hi-C, Zhang et al. (2019), AT | N/A | 409,341,246 | 697,125 |
| Hi-C, Zhang et al. (2019), EG1 | N/A | 400,670,452 | 975,455 |
| Hi-C, Zhang et al. (2019), MG1 | N/A | 362,122,076 | 924,057 |
| Hi-C, Zhang et al. (2019), LG1 | N/A | 445,457,382 | 1,227,761 |

**d** RCMC cell inputs by condition & replicate

| | Replicate 1 | Replicate 2 |
|---|---|---|
| asynchronous | 10M (across 2 tubes) | 10M (across 2 tubes) |
| prometaphase (PM) | 20M (across 4 tubes) | 16M (across 3 tubes) |
| ana/telophase (AT) | 14.5M (across 3 tubes) | 11.5M (across 2 tubes) |
| early G1 (EG1) | 25M (across 5 tubes) | 25M (across 5 tubes) |
| mid G1 (MG1) | 25M (across 5 tubes) | 25M (across 5 tubes) |
| late G1 (LG1) | 25M (across 5 tubes) | 25M (across 5 tubes) |

**e** Reproducibility analysis of M-to-G1 RCMC across conditions and replicates

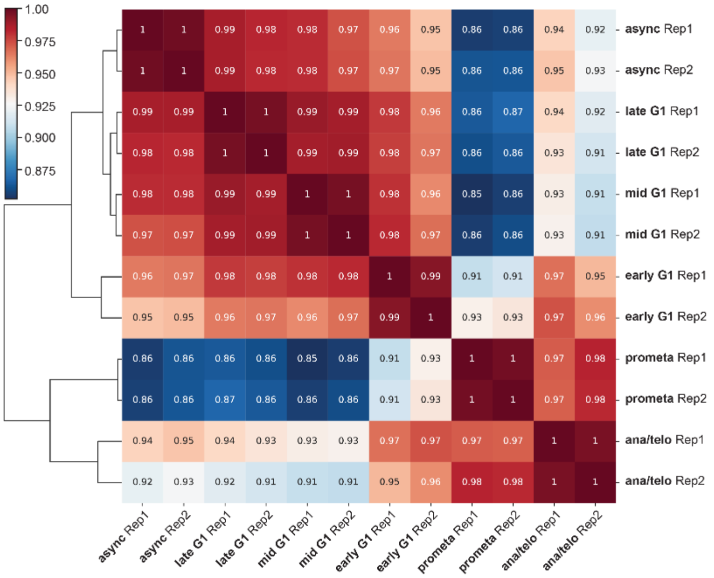


**Supplementary Figure 1. Overview of RCMC data collection, depth, and reproducibility.** (**a**) Experimental workflow for nocodazole arrest-release cell sample collection for RCMC. Flow cytometry plots show the representative gating strategy used to select pure M-to-G1 cell populations based on pMPM2 (prometaphase), mCherry (MD), and DAPI (DNA) signal. (**b**) Summary of the capture efficiency for each of the five regions for which probes were designed. The locations and sizes of the regions, the number of total mapped fragments genome-wide, the number of mapped fragments with at least end within a captured region, and the capture efficiencies are given. (**c**) Table of uniquely mapped RCMC reads genome-wide and within captured ROIs for the five M-to-G1 transition timepoints and the asynchronous conditions. Locus-specific quantifications of unique contacts are shown for the asynchronous condition and reflect ligation products for which both fragment ends are within the captured ROI. A comparison against previously published Hi-C data[1] is also provided. (**d**) Table of M-to-G1 cell sample inputs for RCMC, separated by condition and replicate. Samples were split across multiple tubes treated in parallel containing ~5M cells apiece from MNase digestion onwards and recombined into a single tube during library prep. (**e**) Measurement of reproducibility between M-to-G1 RCMC samples across conditions and replicates. Reproducibility scores are determined using HiCRep[2] at 5 kb resolution, averaged across all five captured loci, and clustered according to similarity.

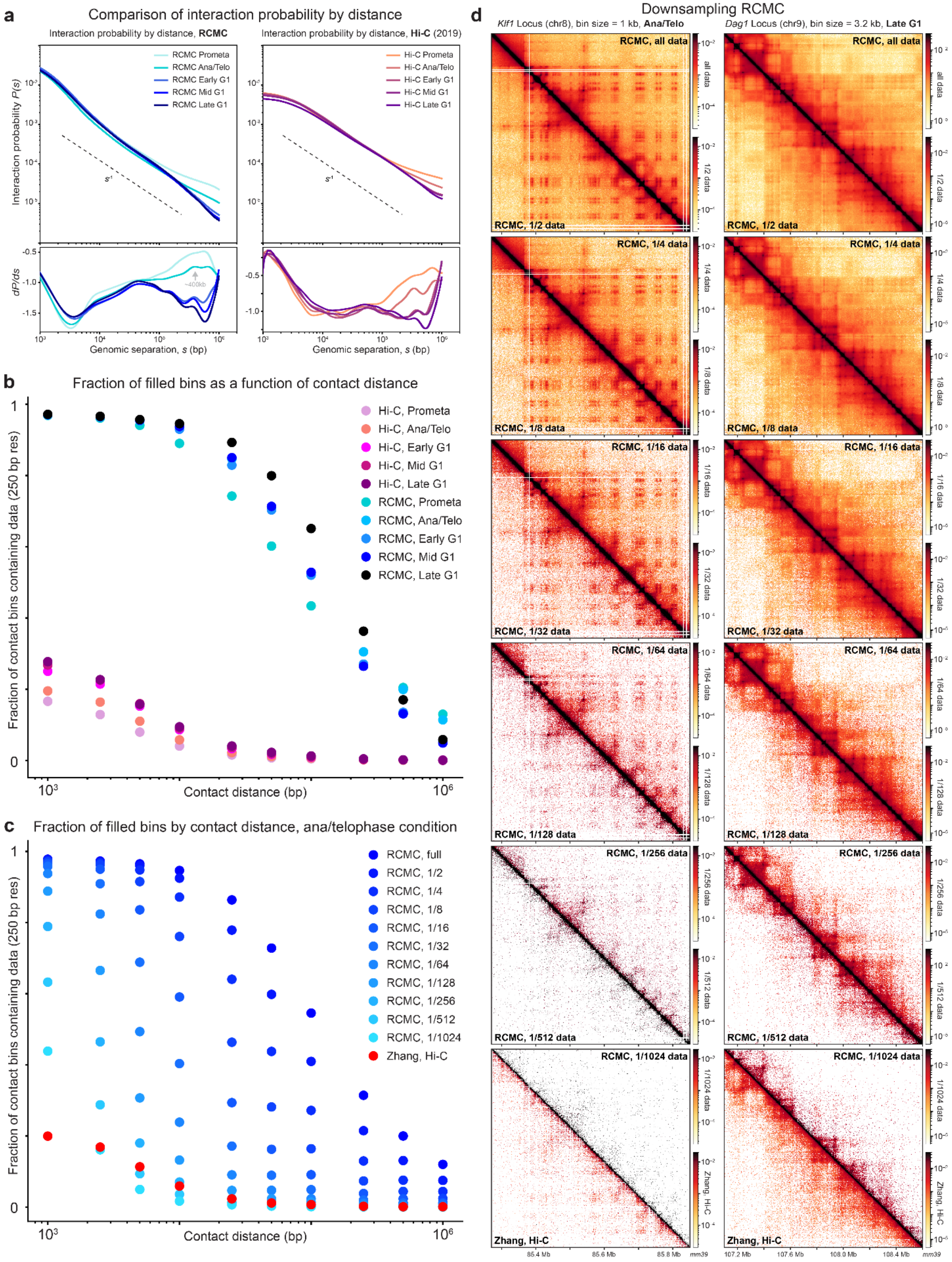


**Supplementary Figure 2. RCMC more deeply and efficiently maps target loci than Hi-C.** (**a**) Interaction probability curves comparing the interaction frequency at different genomic separations ($s$) for the five M-to-G1 conditions, shown for RCMC on the left and Hi-C[1] on the right. The first derivative of these $P(s)$ curves is shown at the bottom. (**b**) Benchmarking comparison of RCMC's ability to fill out high-resolution contact matrices against Hi-C[1]. Region-averaged calculations are shown for both methods across the M-to-G1 datasets. The x-axis shows the contact distance in bp, and the y-axis shows the fraction of all bins at a given contact distance within the captured locus that contain at least one read at 250 bp resolution. (**c**) As in (b), benchmarking comparison of successively downsampled RCMC's ability to fill out high-resolution contact matrices against Hi-C for the ana/telophase condition. Downsampling was applied to all mapped ligated read pairs in increasing powers of 2 until $(1/2)^{10}$, or 1/1024[th], of the initial dataset. (**d**) Contact map comparisons of successively downsampled RCMC data, starting from the full dataset (topmost) down to 1/1024[th] (bottom) and with a comparison against the Hi-C dataset. Maps are shown for the *Klf1* locus at 1 kb resolution in the ana/telophase condition on the left and for the *Dag1* locus at 3.2 kb resolution in the late G1 condition on the right.

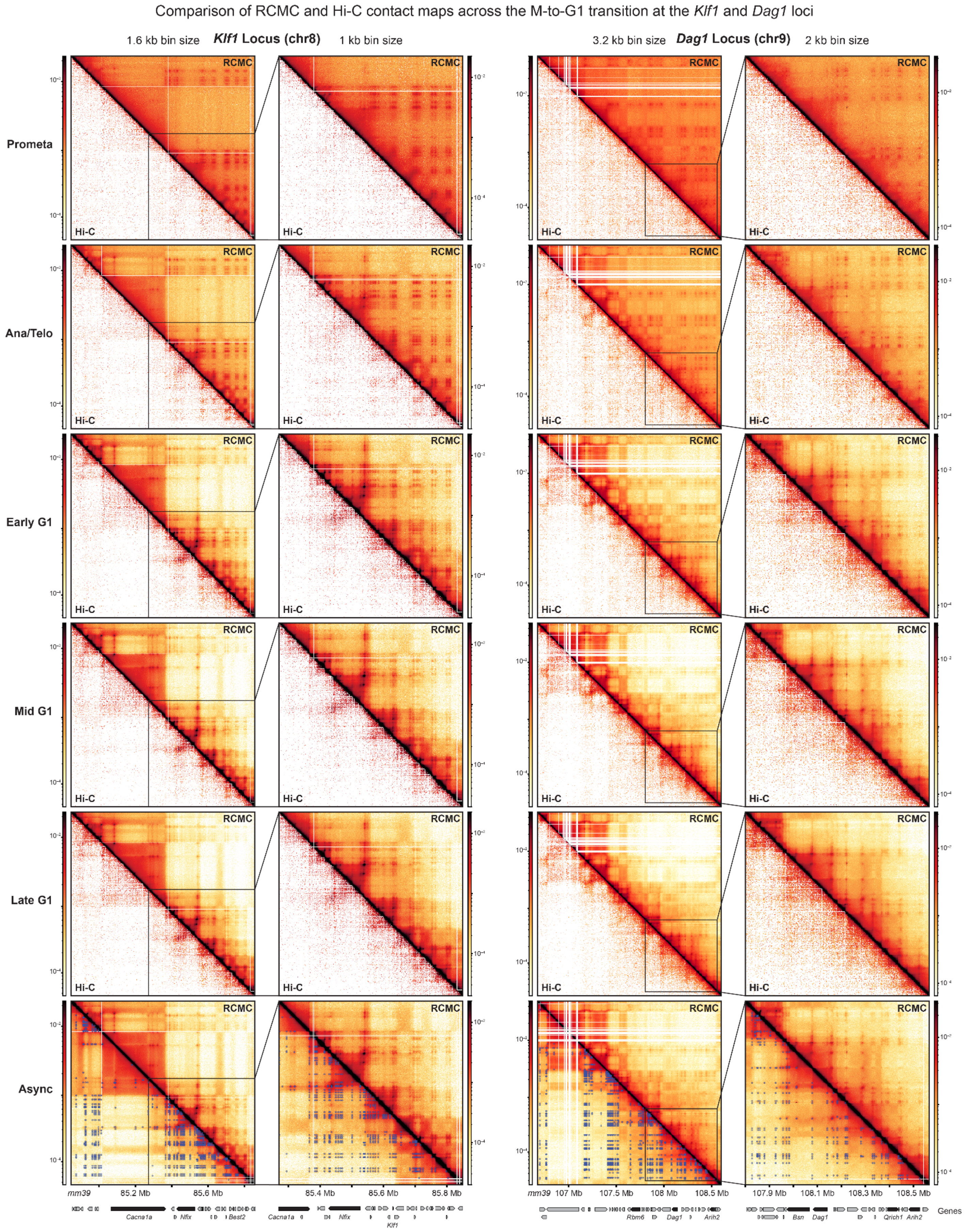


**Supplementary Figure 3. RCMC maps mitotic exit at the *Klf1* and *Dag1* loci.** Contact map comparisons of RCMC against previously generated Hi-C data[1] at the *Klf1* and *Dag1* loci across the M-to-G1 transition. Full capture regions are shown for both loci at 1.6 kb and 3.2 kb resolution, respectively, along with zoom-ins at 1 kb and 2 kb resolution, respectively. The asynchronous RCMC dataset is shown at the bottom, with an overlay of the superset of all annotated interactions below the diagonal. Gene annotations are shown below the contact maps and signal intensity scales are shown next to the maps.

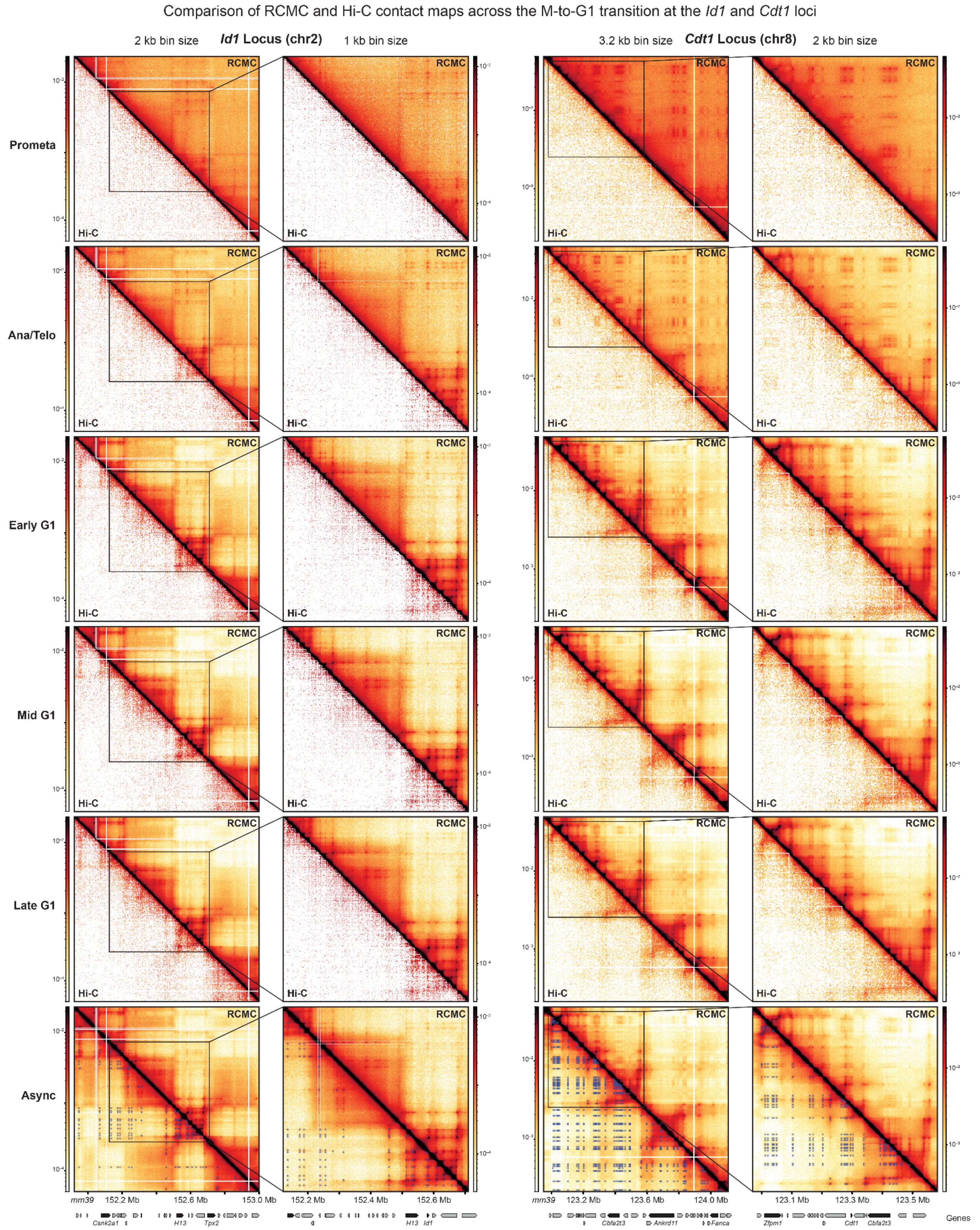


**Supplementary Figure 4. RCMC maps mitotic exit at the *Id1* and *Cdt1* loci.** Contact map comparisons of RCMC against previously generated Hi-C data[1] at the *Id1* and *Cdt1* loci across the M-to-G1 transition. Full capture regions are shown for both loci at 2 kb and 3.2 kb resolution, respectively, along with zoom-ins at 1 kb and 2 kb resolution, respectively. The asynchronous RCMC dataset is shown at the bottom, with an overlay of the superset of all annotated interactions below the diagonal. Gene annotations are shown below the contact maps and signal intensity scales are shown next to the maps.

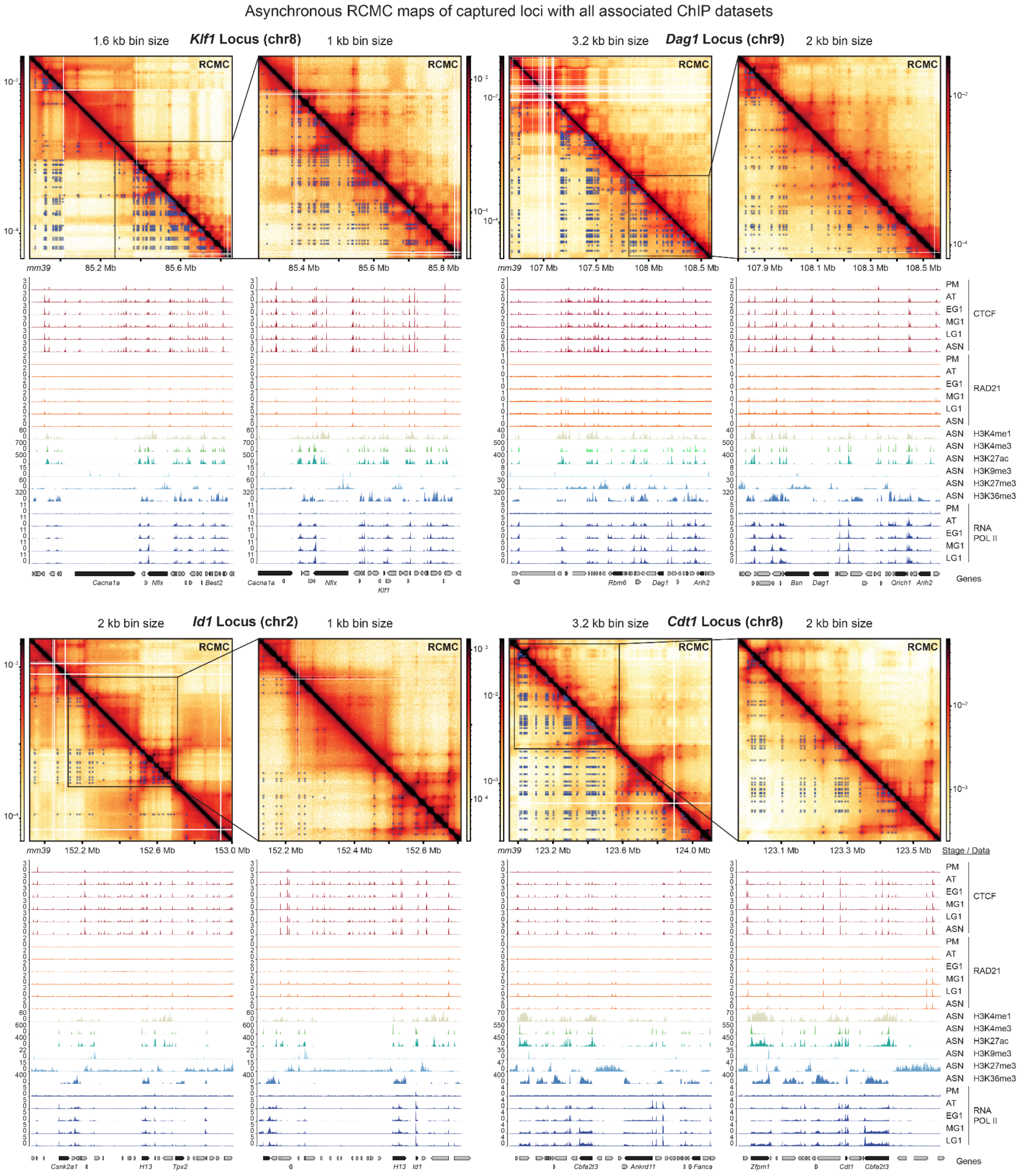


**Supplementary Figure 5. Finely resolved structures in RCMC can be aligned to 1D genomics data.** Asynchronous RCMC contact maps are shown for the *Klf1*, *Dag1*, *Id1*, and *Cdt1* loci alongside zoom-ins at finer resolutions. The superset of annotated interactions is shown below the diagonal. Gene annotations and ChIP-seq tracks (Supplementary Table 1) are shown below the contact maps, while signal intensity scales are shown next to the maps. ChIP-seq tracks include condition-separated CTCF, RAD21, and RNA Pol2 datasets and asynchronous H3K4me1, H3K4me3, H3K9me3, H3K27me3, and H3K36me3 datasets.

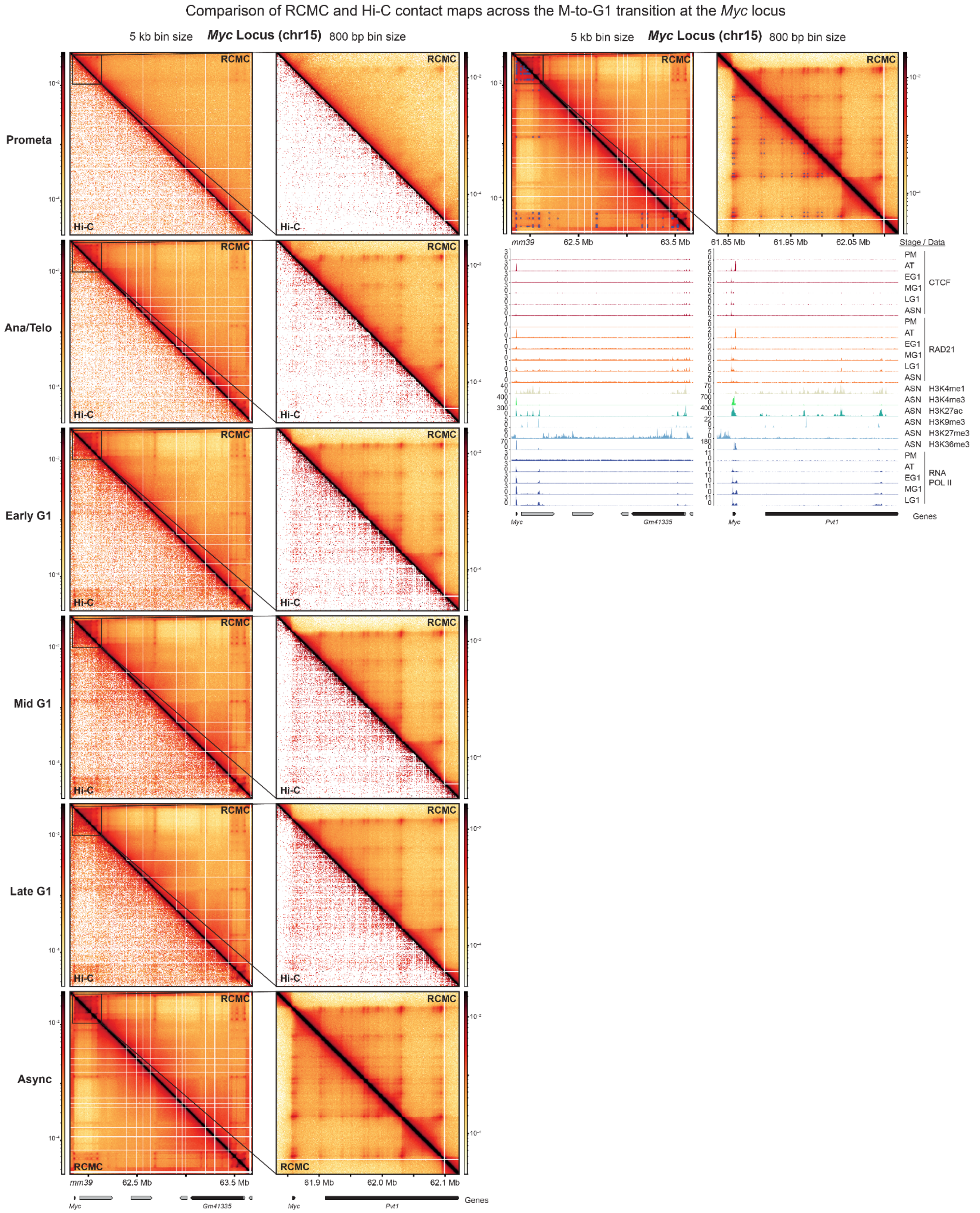


**Supplementary Figure 6. RCMC maps mitotic exit at the *Myc* locus.** (Left) Contact map comparisons of RCMC against previously generated Hi-C data[1] at the *Myc* locus across the M-to-G1 transition. The full capture region is shown at 5 kb resolution along with a zoom-in at 800 bp resolution, and the asynchronous RCMC dataset is shown at the bottom. (Right) Contact maps of the asynchronous dataset are shown with an overlay of the superset of all annotated interactions below the diagonal. Gene annotations are shown below the contact maps and ChIP-seq tracks (Supplementary Table 1) are shown below the contact maps, while signal intensity scales are shown next to the maps.

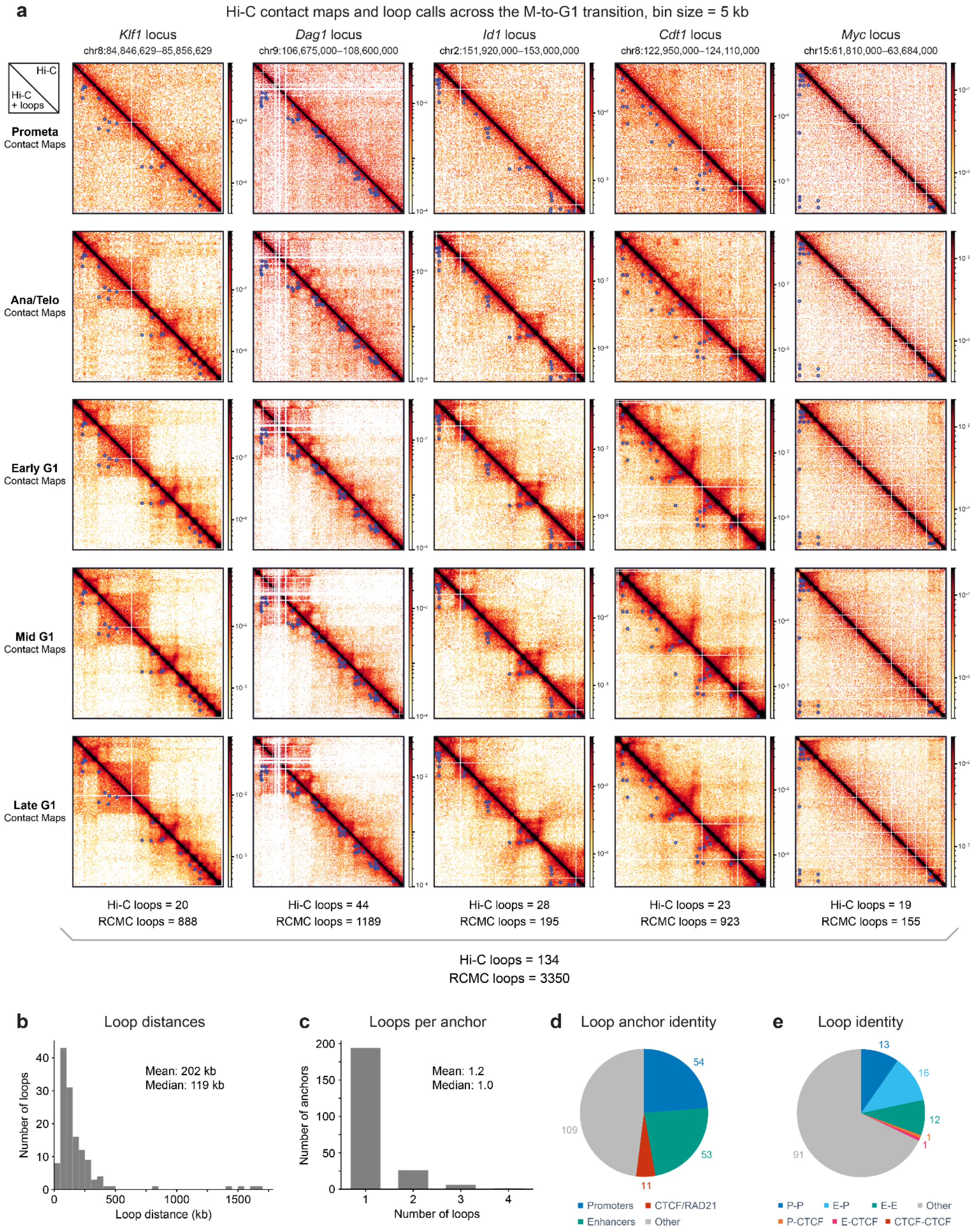


**Supplementary Figure 7. Hi-C detects far fewer loops within the five RCMC regions.** (**a**) Array of contact maps showing previously published Hi-C data[1] and loop calls (below the diagonal) for all M-to-G1 timepoints and capture loci. The superset of M-to-G1 loop calls was generated by merging calls within 10 kb of each other into a single averaged point coordinate. All maps are shown at 5 kb bin size, and the number of loops called in Hi-C and RCMC for each locus is noted below. (**b-c**) Histograms of (**b**) loop interaction distances and (**c**) the number of loops formed by each called anchor. (**d**) Venn diagram of annotated loop anchors by their genomic identity, determined by chromatin features within 1 kb of anchor sites. Promoters were identified as annotated transcription start sites ±2 kb, enhancers as non-promoter regions with overlapping H3K4me1 and H3K27ac ChIP-seq peaks, and CTCF/RAD21 as non-promoter and non-enhancer sites with overlapping CTCF and RAD21 ChIP-seq peaks. Anchors with multiple overlapping genomic features were hierarchically classified into a single classification, with promoters taking precedence, then enhancers, and finally CTCF/RAD21. Anchors designated as “other” do not overlap promoters, enhancers, nor CTCF/RAD21. (**e**) Venn diagram of annotated loops by the genomic identity assigned in (**d**), with P designating promoters, E designating enhancers, and CTCF designating CTCF/RAD21.

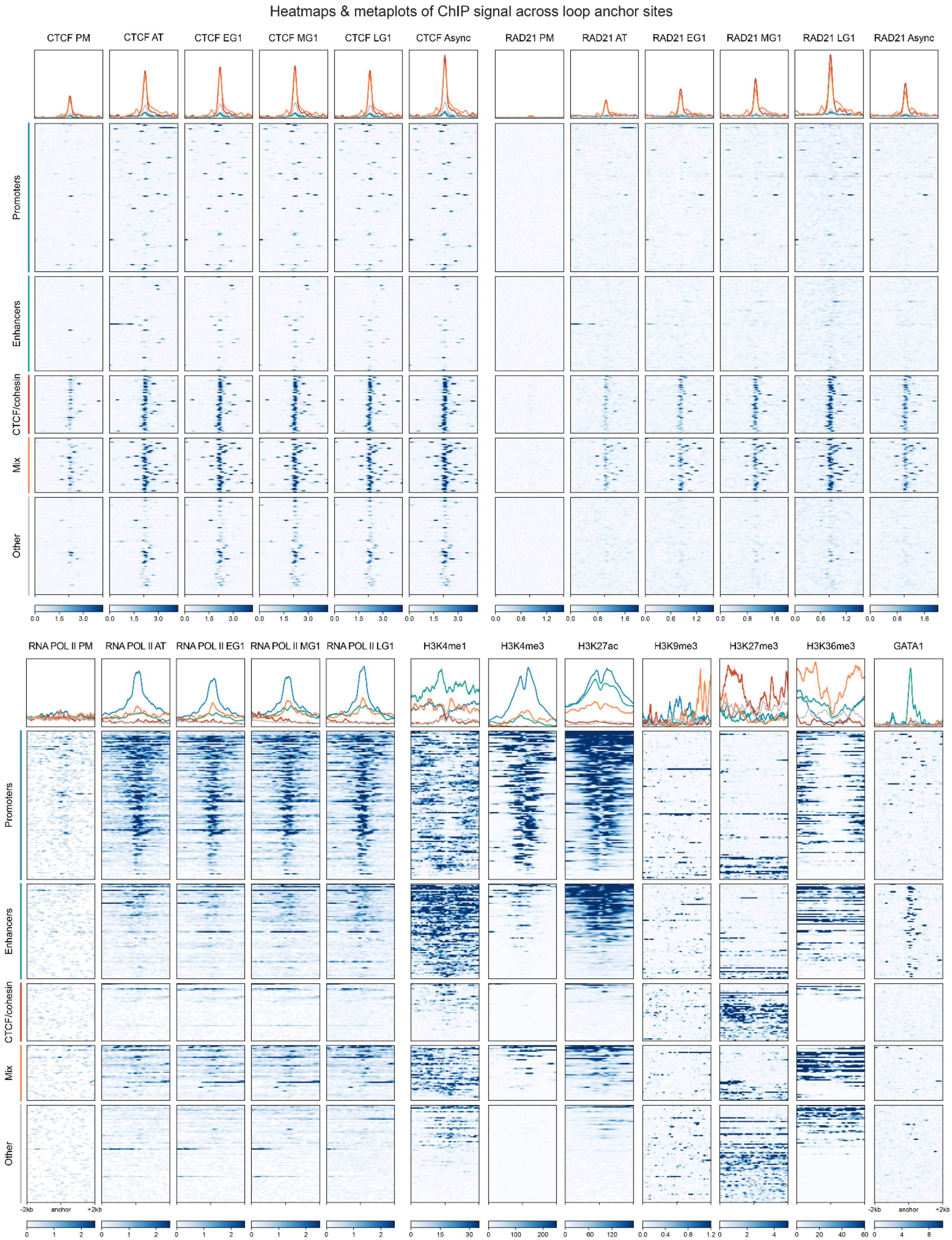


**Supplementary Figure 8 Categories of microcompartment anchors can be defined by their chromatin features.** Metaplots (above) and heatmaps (below) depicting ChIP-seq (Supplementary Table 1) signal at annotated loop anchors as defined in Fig. 2d, including promoters (blue), enhancers (green), CTCF and RAD21-bound (red), a mix of both a cis-regulatory element and CTCF/cohesin (orange), and other (gray). Features are plotted in a 2 kb window centered on the anchor.

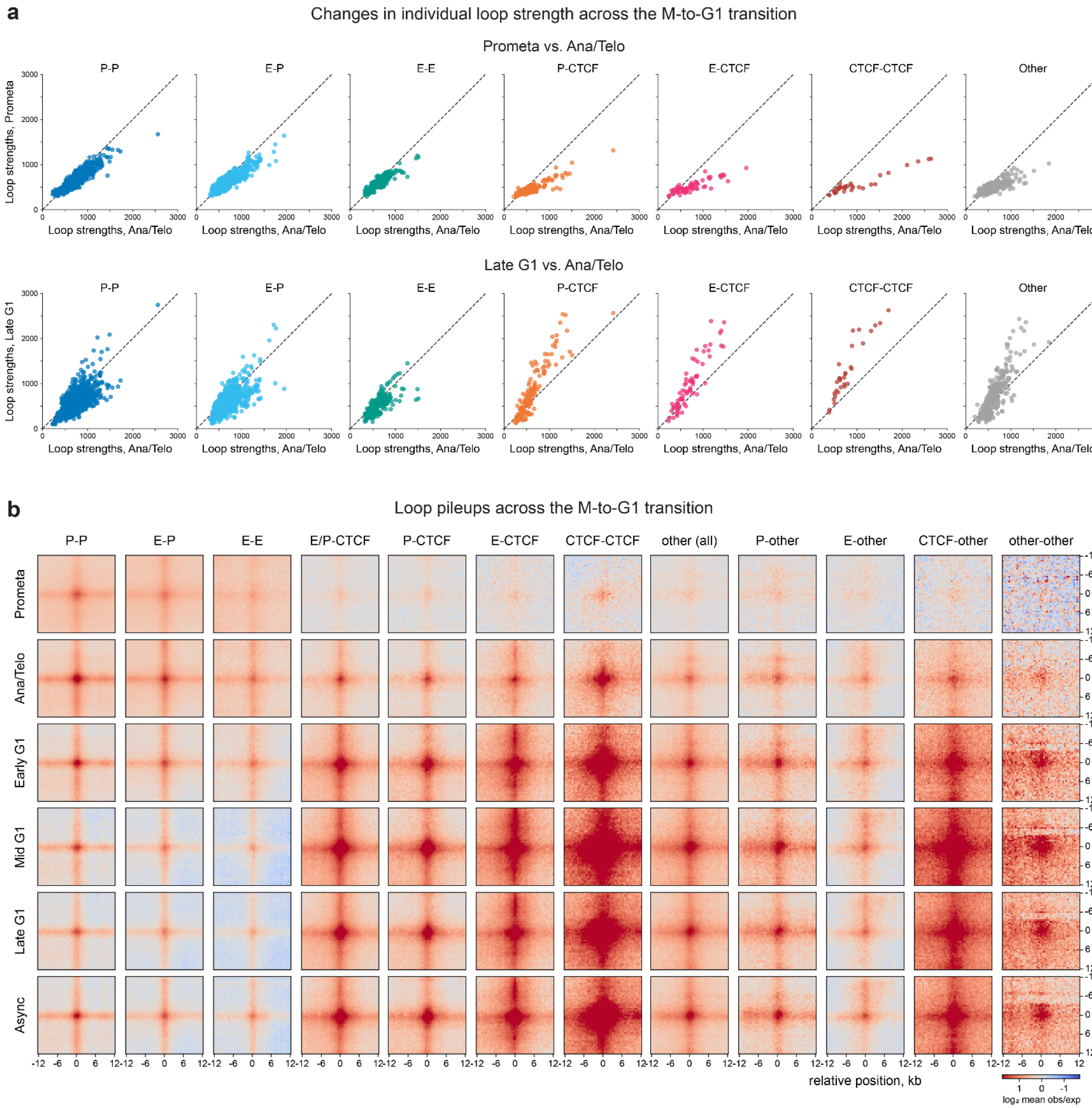


**Supplementary Figure 9. Quantification of M-to-G1 RCMC experiments and loop strengths.** (**a**) Plots of individual loop strengths of each loop category in Fig. 2e for the prometaphase (top) and late G1 (bottom) conditions, plotted against the strengths in the ana/telophase condition (x-axes). Strengths are calculated as the integrated observed loop signal divided by the expected background signal from local *P(s)* curves. The local *P(s)* curves used in this "observed over expected" strength calculation are determined by the loop distance and the dataset's interaction decay curve. These panels show "pure" loops using exclusive loop categorizations, wherein loops anchored by both CREs and CTCF/RAD21 at a single site have been removed. (**b**) Expanded array of the aggregate peak analysis (APA) plots shown in Fig. 3b, separated to show various loop classifications across the M-to-G1 transition and for the asynchronous condition. Plots show a 24 kb window centered on the loop at 500 bp resolution, and the loops plotted here and in all subsequent panels follow the "exclusive" definition of loop identity as in 2g (CRE sites do not overlap with CTCF).

Comparison of RCMC contact maps across condensin depletion at the *Klf1* and *Dag1* loci

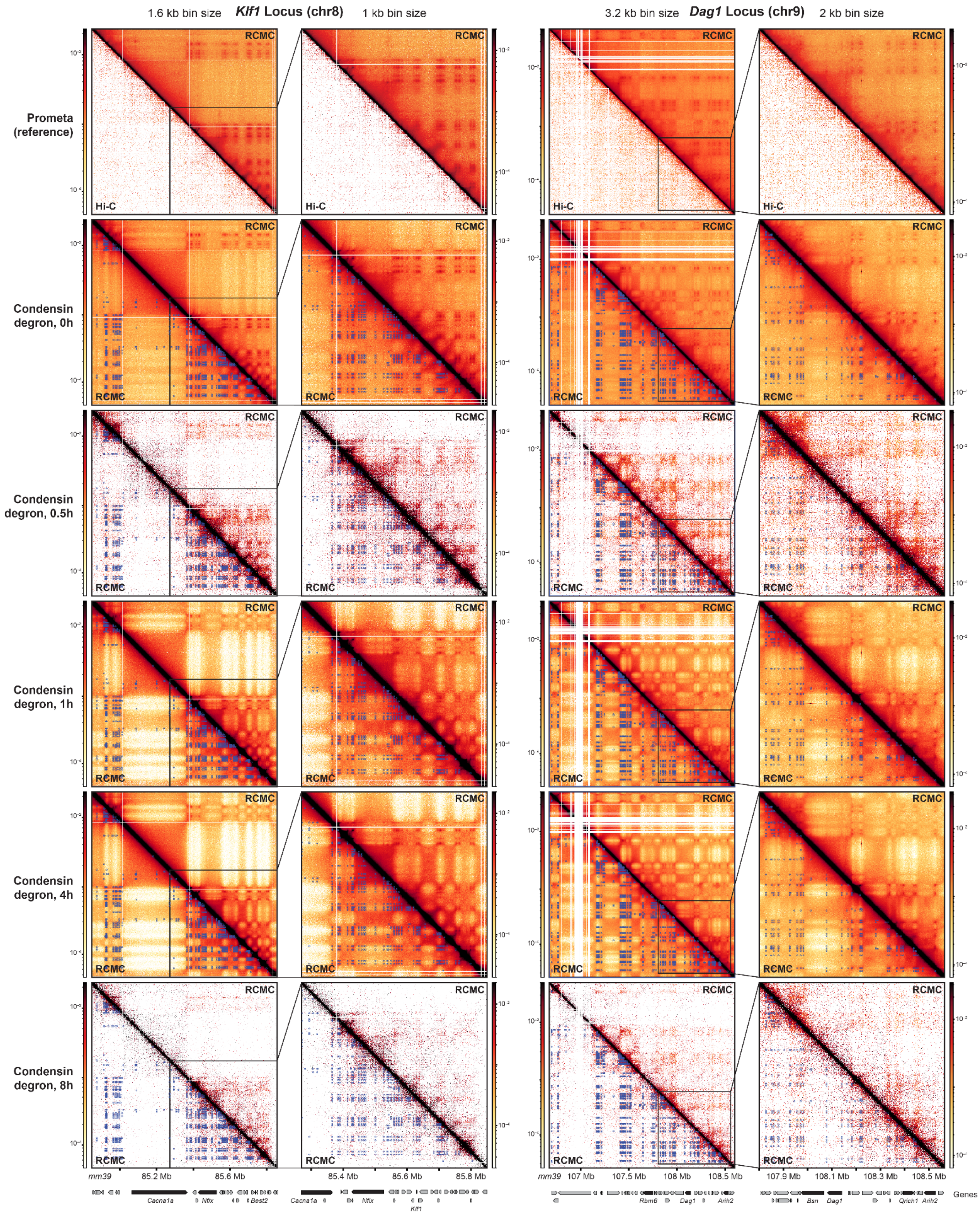


**Supplementary Figure 10. RCMC maps condensin depletion at the *Klf1* and *Dag1* loci.** RCMC contact map comparisons of increasing condensin degradation in prometaphase-arrested cells at the *Klf1* and *Dag1* loci. Condensin degradation of nocodazole-treated SMC2-mAID cells was induced using auxin treatments of 0.5h, 1h, 4h, and 8h, compared against a no treatment (0h) control. Input cell material for RCMC for the 0h, 1h, and 4h conditions was significantly greater than for the 0.5h and 8h conditions. Full capture regions are shown for both loci at 1.6 kb and 3.2 kb resolution, respectively, along with zoom-ins at 1 kb and 2 kb resolution, respectively. The prometaphase M-to-G1 RCMC dataset is shown at the top against previously generated Hi-C data[1] for reference. Gene annotations are shown below the contact maps and signal intensity scales are shown next to the maps.

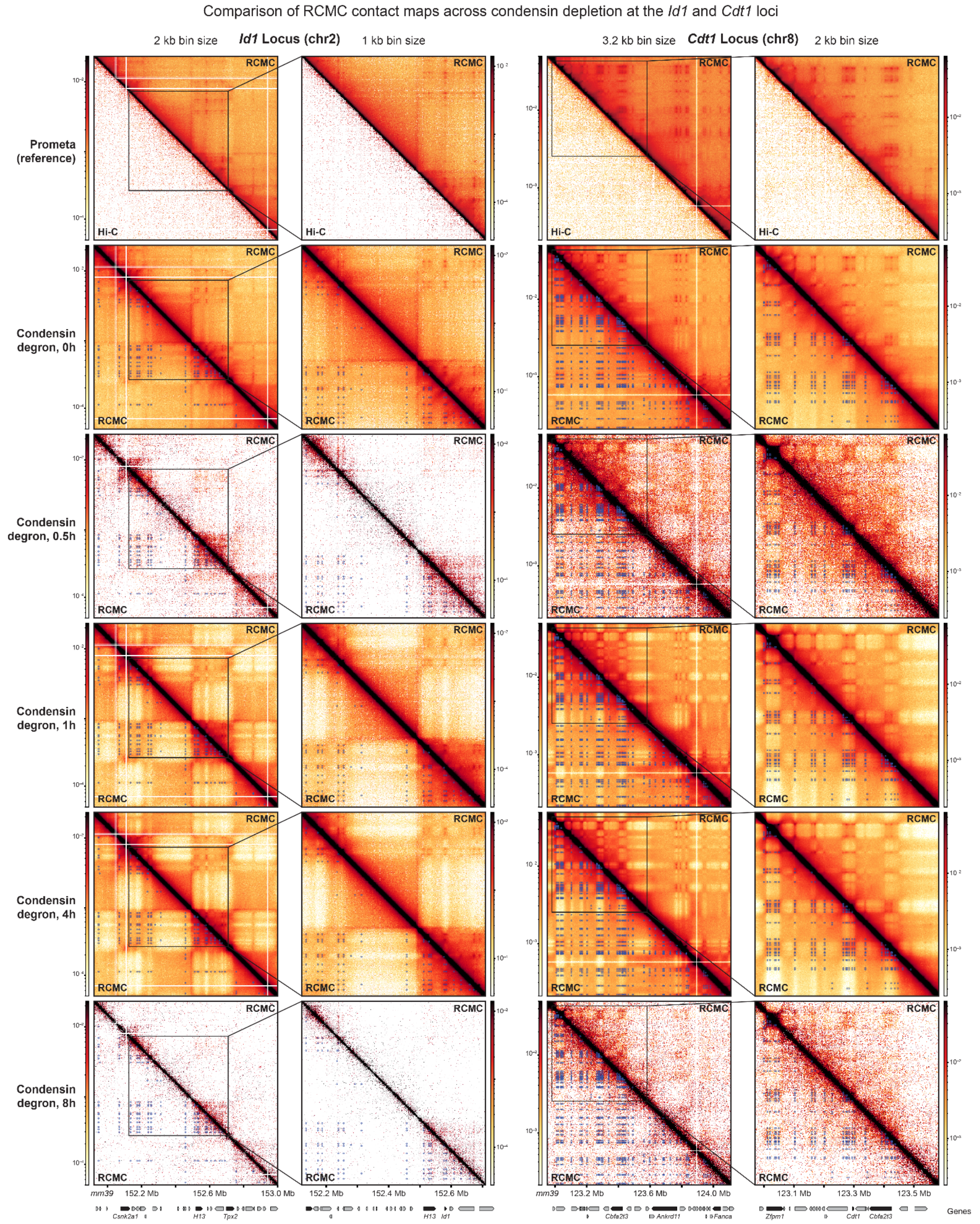


**Supplementary Figure 11. RCMC maps condensin depletion at the *Id1* and *Cdt1* loci.** RCMC contact map comparisons of increasing condensin degradation in prometaphase-arrested cells at the *Id1* and *Cdt1* loci. Condensin degradation of nocodazole-treated SMC2-mAID cells was induced using auxin treatments of 0.5h, 1h, 4h, and 8h, compared against a no treatment (0h) control. Input cell material for RCMC for the 0h, 1h, and 4h conditions was significantly greater than for the 0.5h and 8h conditions. Full capture regions are shown for both loci at 2 kb and 3.2 kb resolution, respectively, along with zoom-ins at 1 kb and 2 kb resolution, respectively. The prometaphase M-to-G1 RCMC dataset is shown at the top against previously generated Hi-C data[1] for reference. Gene annotations are shown below the contact maps and signal intensity scales are shown next to the maps.

Comparison of RCMC contact maps across condensin depletion at the *Myc* locus

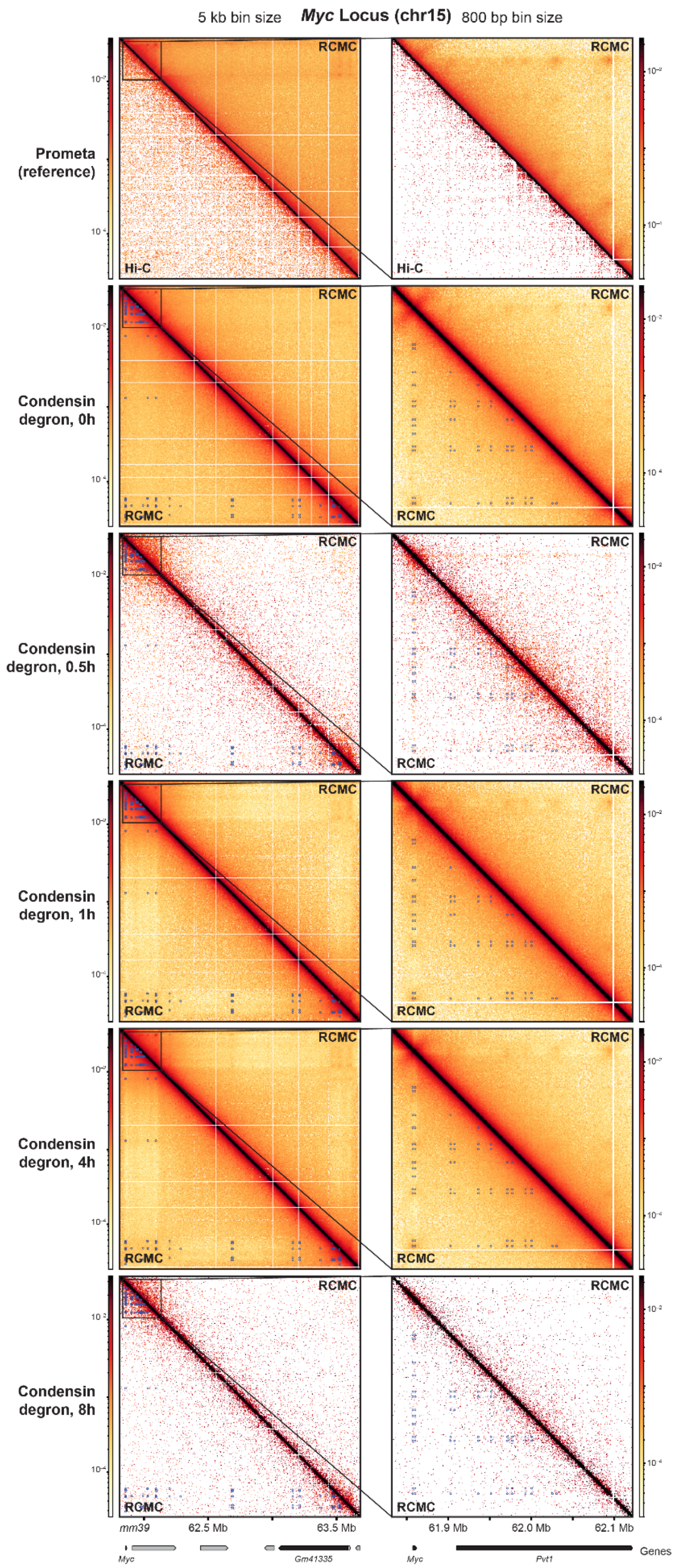


**Supplementary Figure 12. RCMC maps condensin depletion at the *Myc* locus.** RCMC contact map comparisons of increasing condensin degradation in prometaphase-arrested cells at the *Myc* locus. Condensin degradation of nocodazole-treated SMC2-mAID cells was induced using auxin treatments of 0.5h, 1h, 4h, and 8h, compared against a no treatment (0h) control. Input cell material for RCMC for the 0h, 1h, and 4h conditions was significantly greater than for the 0.5h and 8h conditions. The full capture region is shown at 5 kb resolution along with a zoom-in at 800 bp resolution. The prometaphase M-to-G1 RCMC dataset is shown at the top against previously generated Hi-C data[1] for reference. Gene annotations are shown below the contact maps and signal intensity scales are shown next to the maps.

**a** Breakdown of SMC2 depletion sequencing depth

| Read counts comparison | Unique sequencing reads genome-wide | Unique sequencing reads entirely in ROIs |
|---|---|---|
| RCMC, SMC2 depletion, **0h** | 656,234,473 | 191,662,327 |
| RCMC, SMC2 depletion, **0.5h*** | 67,631,171 | 5,609,551 |
| RCMC, SMC2 depletion, **1h** | 520,456,679 | 146,929,186 |
| RCMC, SMC2 depletion, **4h** | 598,908,999 | 199,379,034 |
| RCMC, SMC2 depletion, **8h*** | 34,862,279 | 1,988,799 |

**b** SMC2 depletion RCMC cell inputs

| Cell inputs by replicate | Rep1 | Rep2 | Rep3 | Rep4 | **Total** |
|---|---|---|---|---|---|
| RCMC, SMC2 depletion, **0h** | - | 1.7M | 6.9M | 5.2M | 13.8M |
| RCMC, SMC2 depletion, **0.5h** | 1.7M | 2.4M | - | - | 4.1M |
| RCMC, SMC2 depletion, **1h** | 1.2M | 2.5M | 5.0M | 4.2M | 12.9M |
| RCMC, SMC2 depletion, **4h** | 0.2M | 2.1M | 2.1M | 4.7M | 9.1M |
| RCMC, SMC2 depletion, **8h** | 0.6M | 1.6M | - | - | 2.2M |

**c** Changes in individual loop strength across condensin degradation

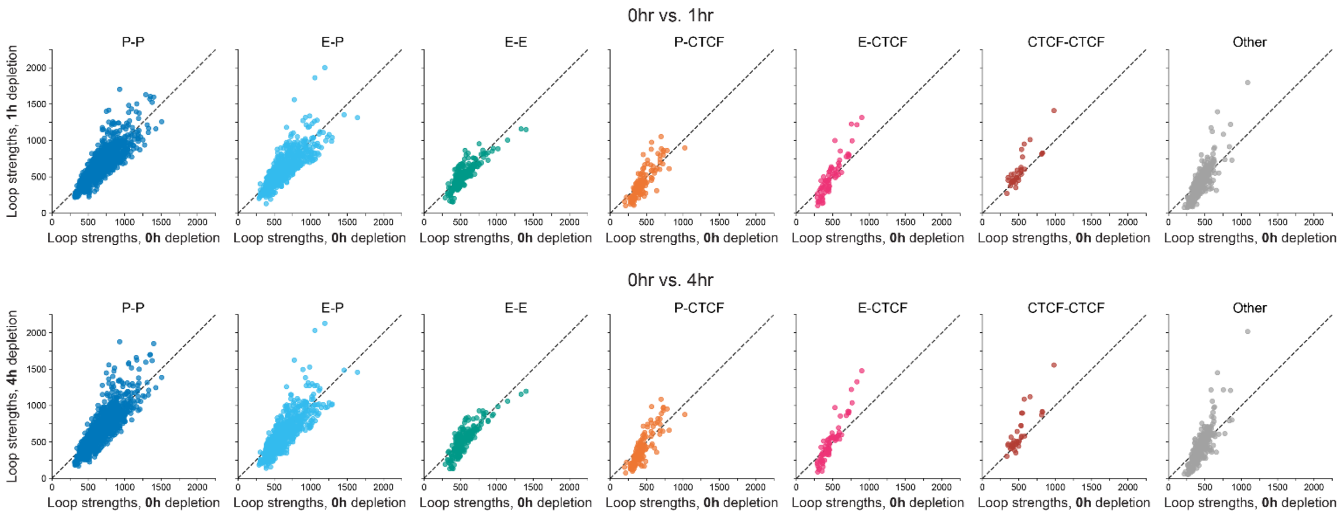


**d** Loop pileups across SMC2 depletion

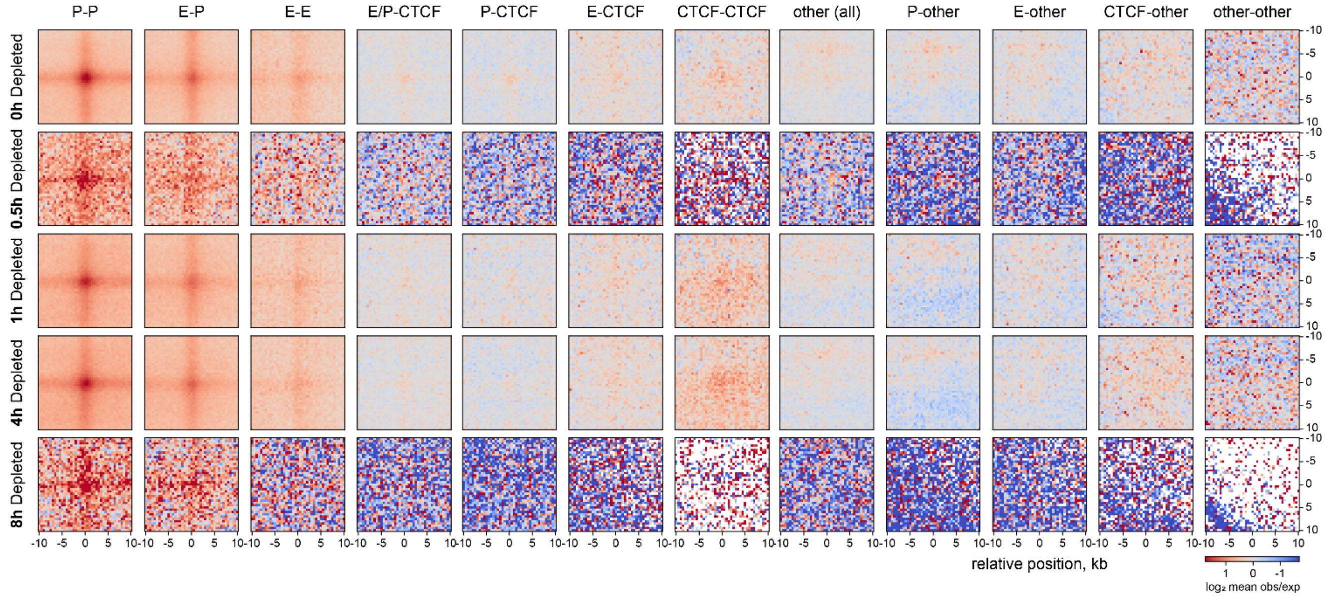


**Supplementary Figure 13. Quantification of SMC2 depletion RCMC experiments and loop strengths.** (**a**) Table of uniquely mapped RCMC reads genome-wide and within captured ROIs for the five condensin depletion timepoints. Asterisks for the 0.5h and 8h conditions denote much shallower sequencing depths resulting from lower cell inputs. (**b**) Table of condensin-depleted cell sample inputs for RCMC, separated by condition and replicate. (**c**) Plots of individual loop strengths of each loop category in Fig. 2e for the 1h (top) and 4h (bottom) condensin depletion conditions, plotted against the strengths in the 0h condition (x-axes). Strengths are calculated as the integrated observed loop signal divided by the expected background signal from local *P(s)* curves. The local *P(s)* curves used in this "observed over expected" strength calculation are determined by the loop distance and the dataset's interaction decay curve. These panels show "pure" loops using exclusive loop categorizations, wherein loops anchored by both CREs and CTCF/RAD21 at a single site have been removed. (**d**) Expanded array of the aggregate peak analysis (APA) plots shown in Fig. 4d, separated to show various loop classifications across condensin depletion. Plots show a 20 kb window centered on the loop at 500 bp resolution, and the loops plotted here and in all subsequent panels follow the "exclusive" definition of loop identity as in 2g (CRE sites do not overlap with CTCF).

**a**

Reproducibility analysis of condensin degron RCMC across conditions and replicates

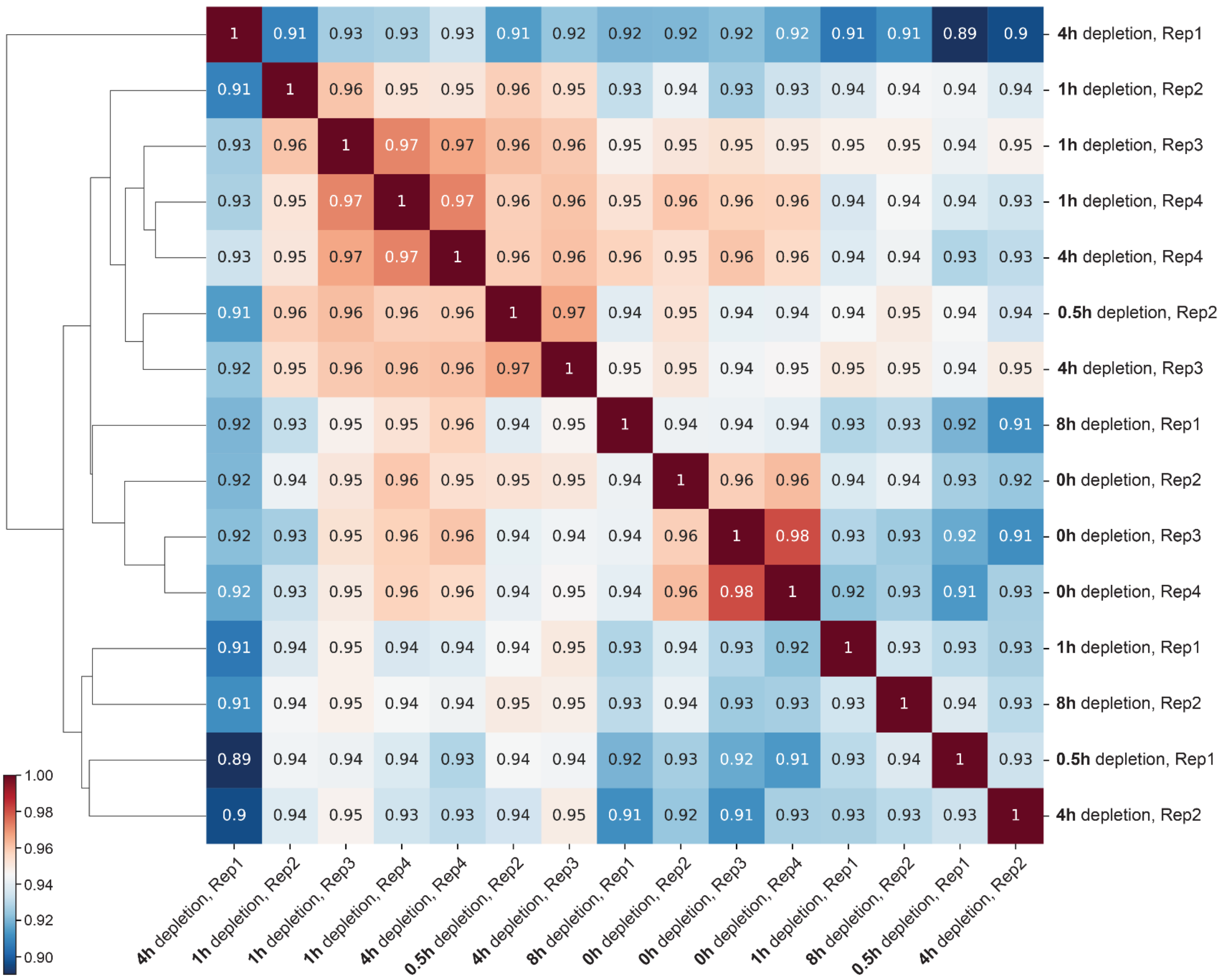


**b**

Unique reads across SMC2 depletion RCMC samples by condition and replicate

| | Rep1 | Rep2 | Rep3 | Rep4 | Total |
|---|---|---|---|---|---|
| RCMC, SMC2 depletion, **0h** | - | 1,063,941 | 113,798,631 | 76,799,755 | 191,662,327 |
| RCMC, SMC2 depletion, **0.5h** | 1,021,022 | 4,588,753 | - | - | 5,609,551 |
| RCMC, SMC2 depletion, **1h** | 633,026 | 5,028,084 | 43,550,381 | 97,717,471 | 146,929,186 |
| RCMC, SMC2 depletion, **4h** | 344,489 | 2,249,551 | 1,837,919 | 194,946,851 | 199,379,034 |
| RCMC, SMC2 depletion, **8h** | 482,762 | 1,506,261 | - | - | 1,988,799 |

>100M
10-100M
1-10M
<1M

**Supplementary Figure 14**. **Quantification of SMC2 degron RCMC experimental reproducibility and replicate depth.** (**a**) Measurement of reproducibility between condensin depletion RCMC samples across conditions and replicates. Reproducibility scores are determined using HiCRep[3] at 5 kb resolution, averaged across all five captured loci, and clustered according to similarity. (**b**) Table of uniquely mapped RCMC reads across captured ROIs for the five condensin depletion timepoints, shown for each replicate. Values are color-coded to highlight magnitudes of difference in sequencing depth across samples, with a legend to the right of the table. Variability in library complexity and sequencing depth stems from the variability in cell inputs shown in Fig. S13b, with higher cellular inputs yielding more complex Micro-C libraries that became larger fractions of the captured RCMC libraries and were ultimately more deeply sequenced.

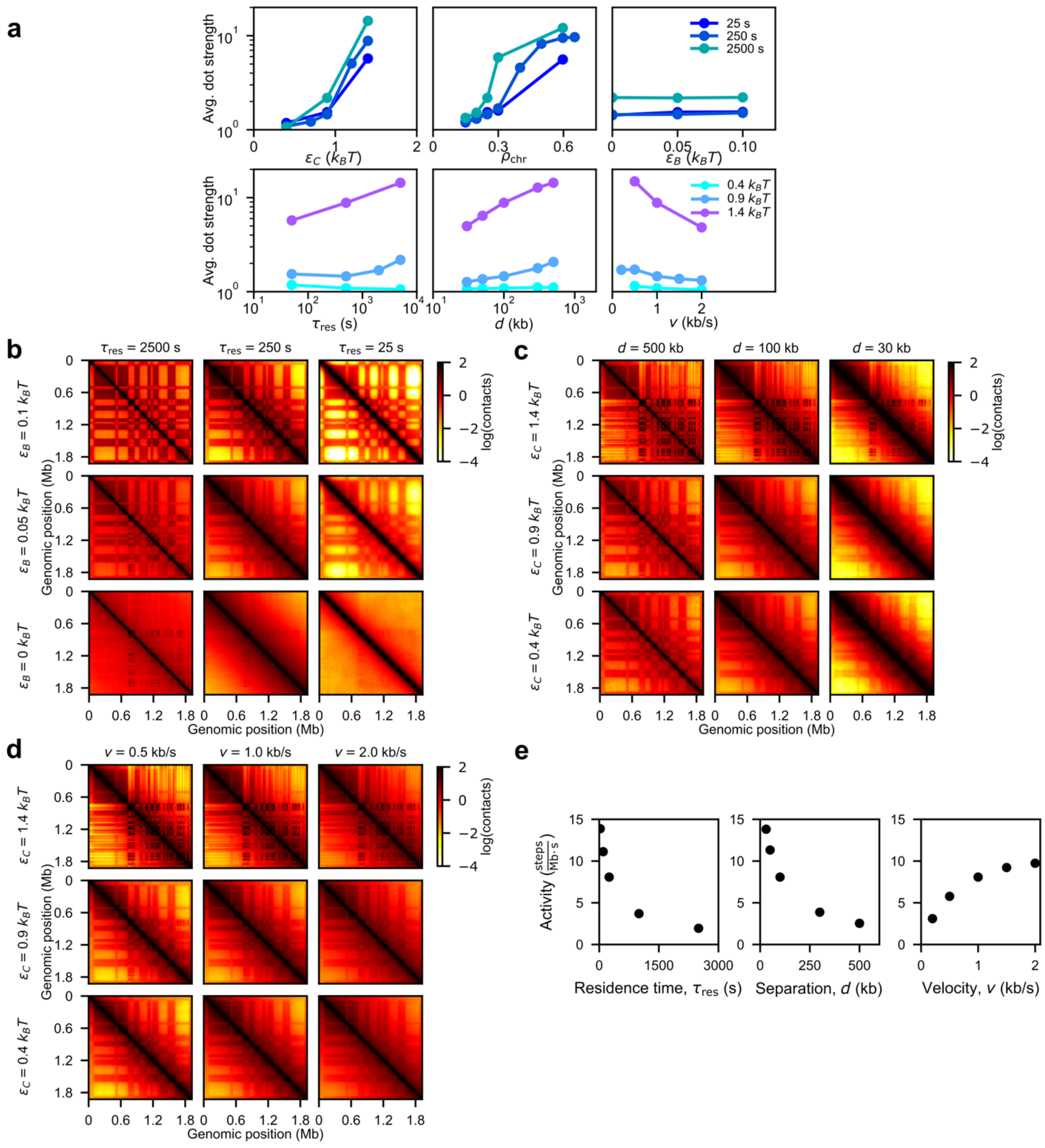


**Supplementary Figure 15. Simulation parameter sweeps of A/B compartment strength, extruder linear density, and extruder velocity.** **(a)** Plots of average dot strengths for different simulation parameters. Top row, from left to right: loop/dot strength versus microcompartment affinity, $\epsilon_C$; chromatin density, $\rho_{chr}$; and A/B compartment affinity, $\epsilon_B$, for different loop extruder residence times, $\tau_{res}$ (different colors). Bottom row, from left to right: dot strength versus loop extruder residence times, $\tau_{res}$; mean separations, $d$, between loop extruders; and loop extrusion velocities, $v$, for different microcompartment affinities, $\epsilon_C$ (different colors). Contact maps from steady-state simulations of the *Dag1* region for different **(b)** loop extruder residence times, $\tau_{res}$ (decreasing from left to right columns) and A/B compartment affinities $\epsilon_B$ (decreasing from top to bottom rows), **(c)** loop extruder linear densities, $1/d$ (increasing from left to right) and microcompartment affinities, $\epsilon_C$ (decreasing from top to bottom), and **(d)** loop extrusion velocities, $v$ (increasing from left to right) and microcompartment affinities, $\epsilon_C$ (decreasing from top to bottom). Velocity is given as the speed of loop growth, i.e., two times the mean translocation speed of each side of a loop extruder. **(e)** Extrusion activity, measured in total extrusion steps per Mb per second, as function of $\tau_{res}$, $d$, or $v$. Standard errors are smaller than the size of the data points.

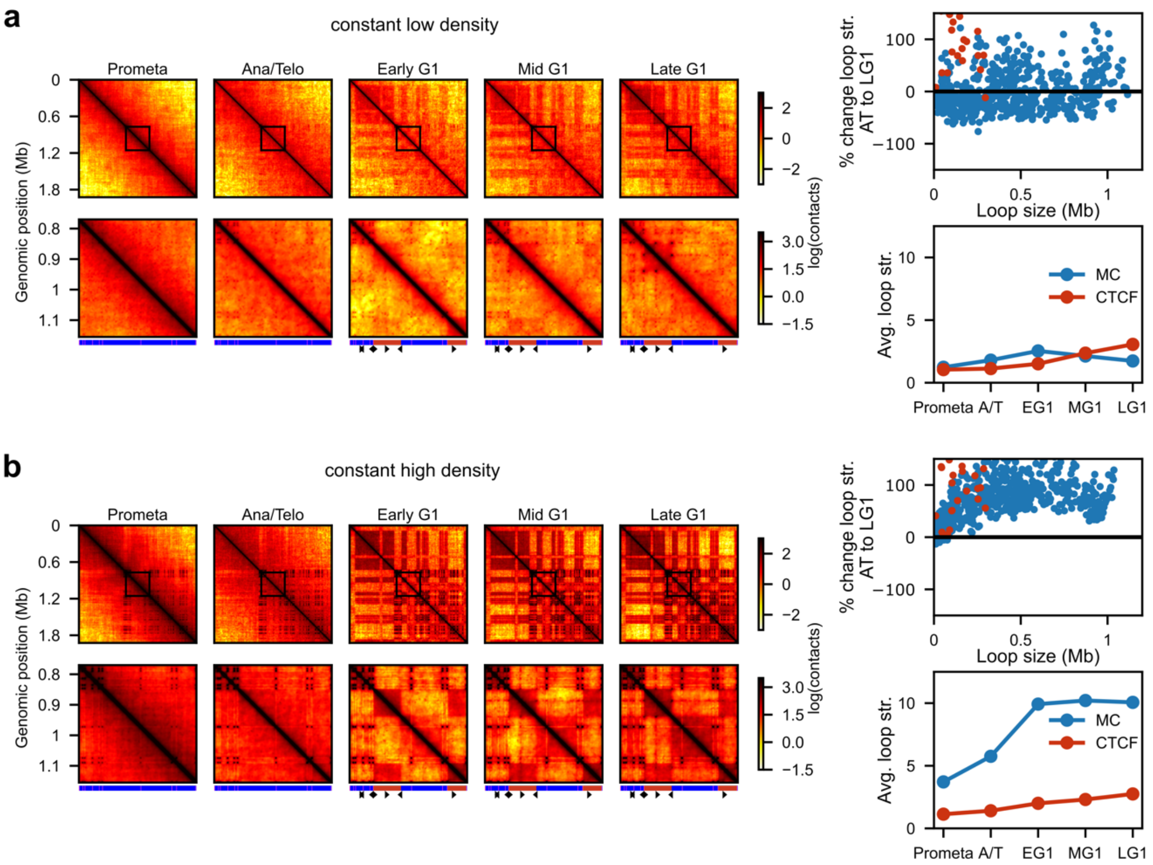


**Supplementary Figure 16. Simulations of the mitosis-to-G1 transition with constant chromatin polymer density.** Results from simulations of the mitosis-to-G1 with density held constant at either **(a)** $\rho_{chr}$=0.25 ("low") or **(b)** $\rho_{chr}$=0.65 ("high"). Left panels show contact maps from various times with the top row showing the full *Dag1* region and the bottom row showing a zoomed-in view of the region marked by the box in the top row. Compartment structure and CTCF sites are indicated below. Right panels show quantification of percent change in loop/dot strength of simulated microcompartments from ana/telophase to late G1 as a function of loop size (top) and average microcompartment and CTCF loop/dot strengths (bottom) throughout the mitosis-to-G1 transition.

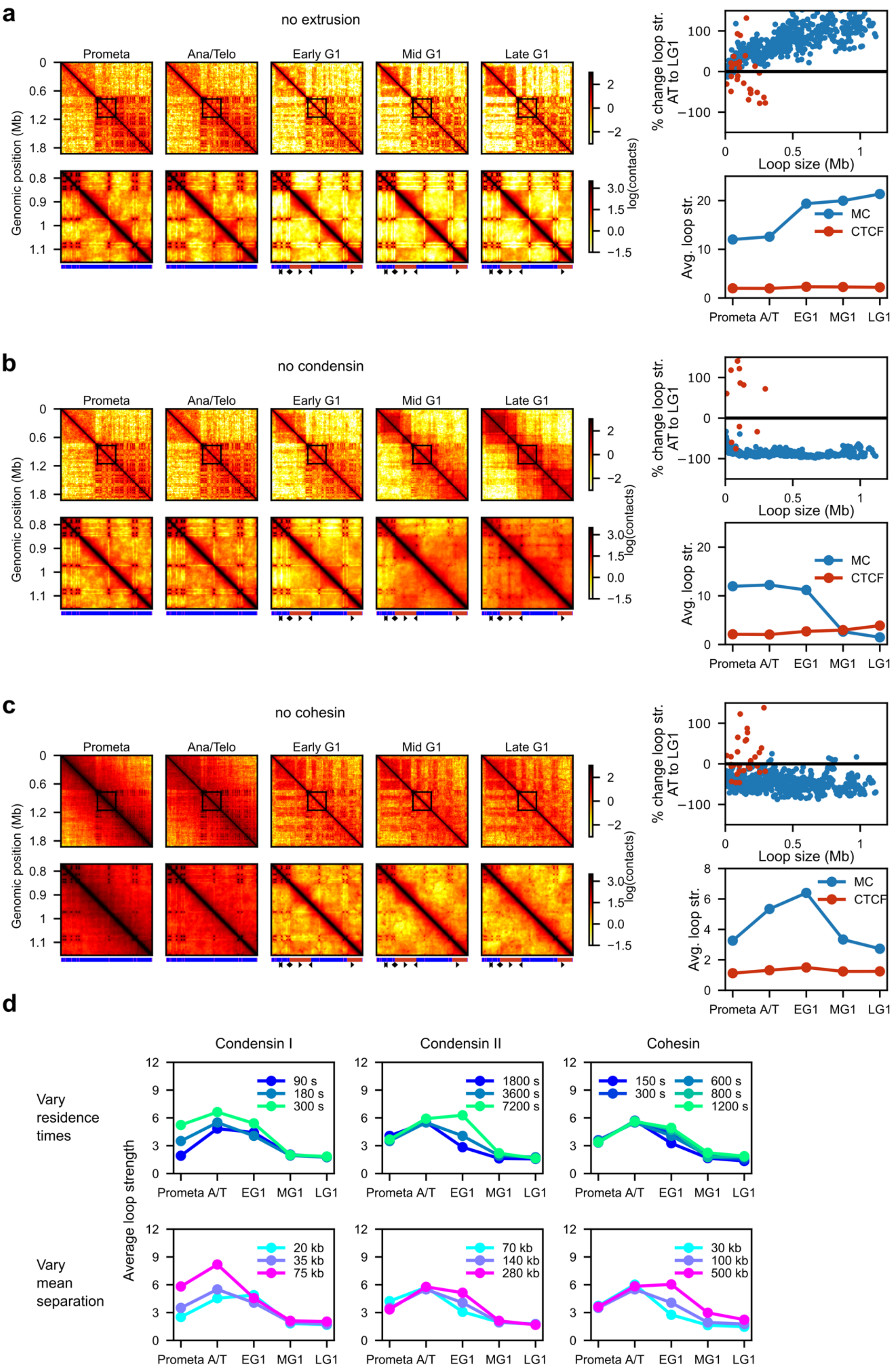


**Supplementary Figure 17. Simulations of the mitosis-to-G1 transition without condensin and/or cohesin and with different loop extrusion parameters.** Results from simulations of the mitosis-to-G1 with **(a)** no loop extruders present, **(b)** no condensin present, but with cohesin in interphase, and **(c)** no cohesin present, but with condensins I and II in prometaphase and ana/telophase. Left panels show contact maps from various times with the top row showing the full *Dag1* region and the bottom row showing a zoomed-in view of the region marked by the box in the top row. Compartment structure and CTCF sites are indicated below. Right panels show quantification of percent change in loop/dot strength of simulated microcompartments from ana/telophase to late G1 as a function of loop size (top) and average microcompartment and CTCF loop/dot strengths (bottom) throughout the mitosis-to-G1 transition. **(d)** Plots of average microcompartment loop/dot strengths throughout the mitosis-to-G1 transition simulations for different loop extrusion parameters. Top row shows loop/dot strengths for different residence times, $\tau_{\text{res}}$, and bottom row shows dot strengths for different mean separations, *d*. Different columns show loop/dot strengths for alterations to different loop extruders with parameters for other loop extruders held fixed.

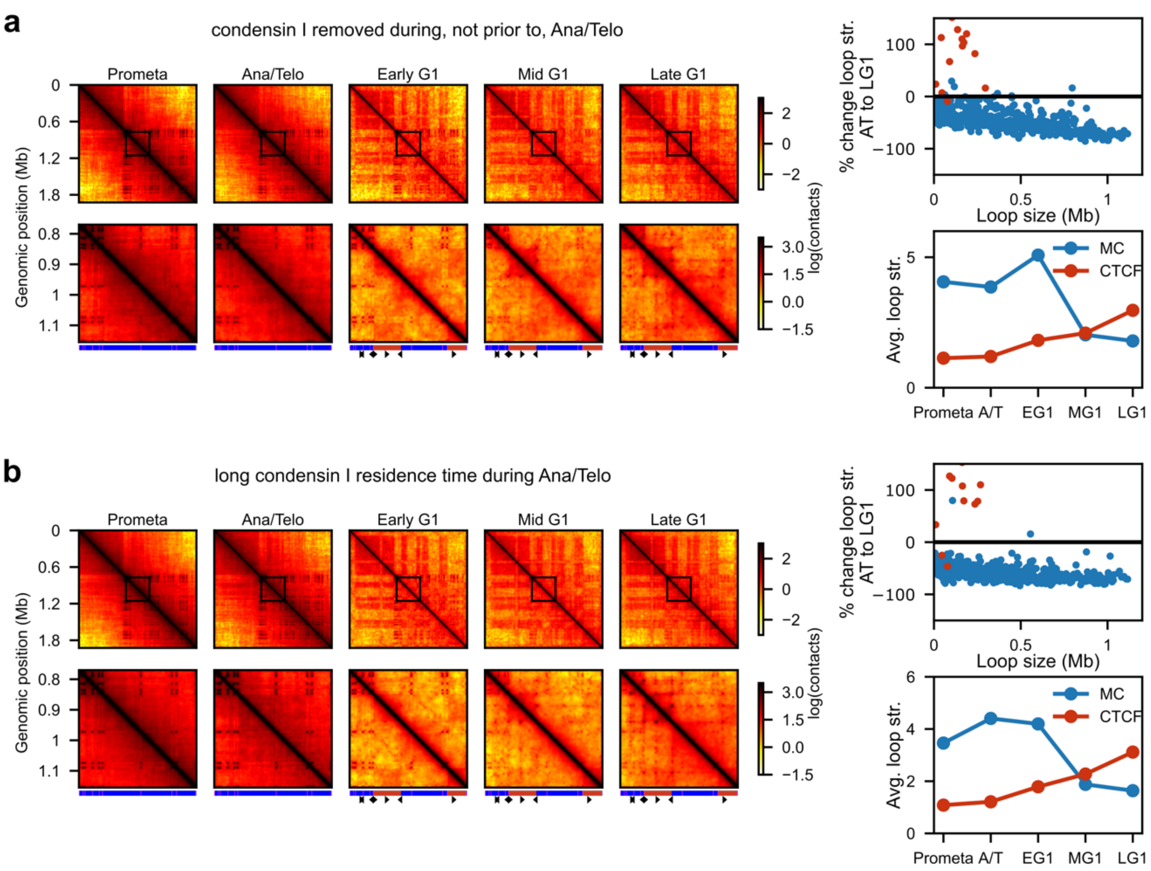


**Supplementary Figure 18. Simulations of the mitosis-to-G1 transition with alterations to condensin I removal timing or ana/telophase turnover.** Results from simulations of the mitosis-to-G1 with **(a)** condensin I removal during ana/telophase, rather than prior to ana/telophase as in the main text, and **(b)** condensin I removal during ana/telophase with residence time of condensin I, $\tau_{res}^{CI}$, increased 10-fold during ana/telophase. Left panels show contact maps from various times with the top row showing the full *Dag1* region and the bottom row showing a zoomed-in view of the region marked by the box in the top row. Compartment structure and CTCF sites are indicated below. Right panels show quantification of percent change in loop/dot strength of simulated microcompartments from ana/telophase to late G1 as a function of loop size (top) and average microcompartment and CTCF loop/dot strengths (bottom) throughout the mitosis-to-G1 transition.

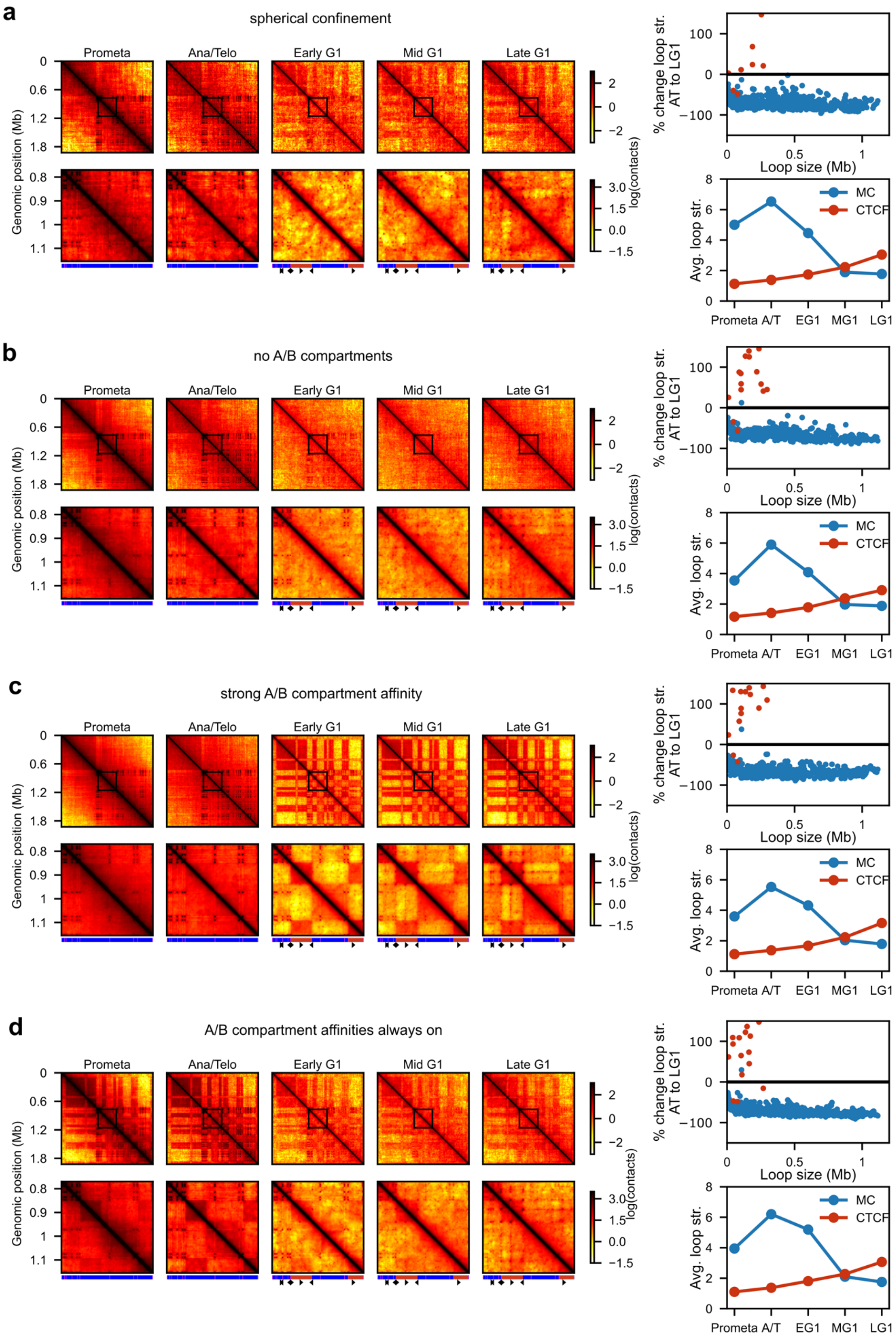


**Supplementary Figure 19. Simulations of the mitosis-to-G1 transition with different boundary conditions or A/B compartment affinities.** Results from simulations of the mitosis-to-G1 with **(a)** no cylindrical boundary condition during mitosis, i.e., only spherical confinement, such that while chromosome density changes, chromosome shape does not; **(b)** no A/B compartment affinity interactions; **(c)** A/B compartment affinity doubled to $\epsilon_B$=0.1 $k_BT$; and **(d)** A/B compartment affinity always on at the typical value of $\epsilon_B$=0.05 $k_BT$, including during prometaphase and ana/telophase. Left panels show contact maps from various times with the top row showing the full *Dag1* region and the bottom row showing a zoomed-in view of the region marked by the box in the top row. Compartment structure and CTCF sites are indicated below. Right panels show quantification of percent change in loop/dot strength of simulated microcompartments from ana/telophase to late G1 as a function of loop size (top) and average microcompartment and CTCF loop/dot strengths (bottom) throughout the mitosis-to-G1 transition.

### MATERIALS AND METHODS

#### *Experimental Procedures*

**Overview of the Region Capture Micro-C experiment**

Region Capture Micro-C (RCMC)[4] was developed by merging Micro-C[5] with tiling Region-Capture of a locus[6,7]. The detailed RCMC protocol is provided as Supplementary Information in the original RCMC publication[4]. Here we summarize the protocol descriptions previously described in Goel *et al.* 2023[4].

The data generated in this manuscript come from merging of multiple replicates. For the mitosis-to-G1 cell cycle synchronized datasets, two RCMC biological replicates were generated for each of the six tested conditions: prometaphase, ana/telophase, early G1, mid G1, late G1, and the asynchronous condition (**Fig. S1d**). For the condensin degron datasets, four RCMC biological replicates were generated across five tested conditions (0h, 0.5h, 1h, 4h, and 8h of depletion) (**Fig. S13a, 13b**). Biological replicates were generated by harvesting (culturing, crosslinking, aliquoting, and snap-freezing) 1-50M cells for each tested condition, after which downstream RCMC steps (Micro-C and Region Capture) were applied to snap-frozen cell aliquots totaling up to 15-25M input cells to generate each biological replicate.

**Cell culture and maintenance**

The G1E-ER4 murine erythroblast cell line was originally gifted by Dr. Mitchel Weiss[8]. Two G1E-ER4 sublines were used in this study: G1E-ER4-mCherry-MD[1], and SMC2-AID-mCherry[3]. G1E-ER4-mCherry-MD cells express the mitotic degradation domain of cyclin B fused to mCherry (used for isolating specific cell populations during the M-G1 transition)[1]. The SMC2-AID-mCherry subline harbors a homozygous, in-frame insertion (auxin-inducible-degron sequence and mCherry) at the SMC2 locus[3]. All lines were maintained under previously described conditions for G1E-ER4 cells[8].

**Synchronization and isolation of mitotic cell populations**

Synchronization and isolation of cell populations during mitotic exit was carried out as previously described[1] with minor modifications (**Fig. 1a, Fig. S1a**). Briefly, actively dividing cells (density: 0.5-0.8 million cells/ml) were treated with 200ng/ml nocodazole for 8.5h for prometaphase arrest. Cells were washed with nocodazole-free media and released for the following timepoints to enrich for specific populations during the mitosis-G1 transition: 40min (ana/telophase), 1h (early G1), 2h (mid G1) or 4h (late G1). After nocodazole treatment and release, cells were sequentially crosslinked with formaldehyde and DSG. Cells were centrifuged for 5min at 1500 rpm and resuspended in PBS with 1% formaldehyde (1 million cells/mL) and incubated with gentle rocking at room temperature for 10 min. Formaldehyde crosslinking was quenched with 0.375M Tris pH 7.5 (room temperature, 5 min). Cells were washed twice with cold PBS and resuspended in 3mM DSG (ProteoChem, c1104-1gm). DSG crosslinking was carried out for 45 min at room temperature with gentle rocking. DSG crosslinking was quenched with 0.375M Tris pH 7.5 (room temperature, 5min). Cells were then permeabilized with 0.1% TritonX-100 and stained with mitosis specific anti-pMPM2 antibody (Millipore, 05-368), for 50 min at room temperature (0.5uL/10 million cells). Secondary antibody staining was carried out for 30 min at room temperature with APC-conjugated F(ab')2-Goat anti-Mouse (Thermo Fisher Scientific, 17-4010-82). Finally, cells were resuspended in FACS buffer with 25ng/mL DAPI and kept on ice. To enrich for populations in specific cell cycle stages during the Mitosis-G1 transition, cells were subjected to flow cytometry sorting using the MoFlo Astrios EQ sorter (Beckman Coulter). The following markers were used to isolate specific cell populations: prometaphase – high mCherry-MD, positive pMPM2, 4N DAPI, ana/telophase- low mCherry-MD, 4N DAPI, G1 populations- negative mCherry-MD, 2N DAPI. Sorted cells were aliquoted and flash frozen in liquid nitrogen.

**SMC2 degradation**

Mitotic depletion of SMC2 was carried out as previously described[3] (**Fig. 4a**). Briefly, G1E-ER4-SMC2-AID-mCherry cells were first arrested in prometaphase with (200ng/mL) nocodazole treatment for either 12h or 15h (for 8h auxin timepoint). During the end of nocodazole treatment, cells were also treated with auxin (1mM) to deplete SMC2 for the following timepoints: 0h, 0.5h, 1h, 4h, and 8h. Total nocodazole treatment time was 12h for all samples except for the 8h auxin treatment which had a total nocodazole treatment time of 15h. Cells were serially crosslinked as described above with 1% formaldehyde and subsequently 3mM DSG. Cells were permeabilized and stained with pMPM2 primary antibody and APC-conjugated F(ab')2-Goat anti-Mouse (Thermo Fisher Scientific, 17-4010-82) as described above. Cells were subjected to flow cytometry to enrich for prometaphase-arrested samples. All samples were sorted for pMPM2+ cells; auxin-treated cells were sorted based on low mCherry signal (indicative of SMC2 degradation).

**Crosslinking**

Crosslinking was performed precisely as in Goel *et al.* 2023[4]. Briefly, cells were doubly crosslinked using 3 mM DSG (ThermoFisher #20593) and 1% formaldehyde (ThermoFisher #28906). Single cells were counted, washed in PBS, and then resuspended in 3mM DSG in PBS at a concentration of 1M cells per mL, mixed gently for 35 min at room temperature after which formaldehyde was added to a final concentration of 1% and gently mixed at room temperature for an additional 10 minutes and finally quenched with Tris buffer pH = 7.5 (K-D Medical #RGE-3370) at a final concentration of 0.375 M. Crosslinked cells were washed twice with 1X PBS, re-counted, and then partitioned into 1-5M cell aliquots that were pelleted and snap-frozen in liquid nitrogen for storage at –80°C.

**Micrococcal nuclease (MNase) titration**

MNase titration was performed as described in Goel *et al.* 2023. Ideal digestion conditions were determined for each batch of crosslinked cells by treating 1M cell samples with varying amounts of MNase and digesting at 37°C for 20 min on a thermomixer. Digested chromatin underwent crosslink reversal, DNA purification, and gel-based separation to visualize the fragment size distribution. Ideal digestion concentrations were identified by samples digested to primarily (~80%) monomeric fragments (150-200 bp), few (~15-20%) dimeric fragments (250-350 bp), and a faint but visible band (<5%) of trimeric fragments (400-500 bp).

**Sample inputs**

RCMC was performed on cellular inputs ranging from 1-25M of total post-crosslinking sample for each sample condition and replicate. For samples with ample crosslinked material, multiple 5M cell samples were carried in parallel through the protocol to maximize library complexity and combined into a single tube during library preparation. For scarcer samples (e.g., 5M cells or less), RCMC was performed on all available material in a single tube. All M-to-G1 RCMC datasets in this manuscript were generated from two biological replicates of 15-25M input cells each. The condensin degron RCMC datasets in this manuscript were generated from 2-4 biological replicates of 0.2-5M input cells each, with significantly less crosslinked material available for the 0.5h and 8h auxin treatment conditions than for the 0h, 1h, and 4h conditions.

**Micrococcal nuclease digestion**

As described in Goel *et al.* 2023, cell membranes were solubilized to extract intact nuclei by resuspending crosslinked 5M cell pellets in Micro-C Buffer #1 (MB#1; 50 mM NaCl, 10 mM Tris-HCl pH = 7.5, 5 mM MgCl2, 1M CaCl2, 0.2% NP-40 Alternative (Millipore Sigma #492018), 1x Protease Inhibitor Cocktail (Sigma-Aldrich #5056489001)) at 1M cells per 100 µL for 20 min on ice. Following an MB#1 wash, samples were resuspended in 100 µL MB#1 and the ideal amount of 20 U/µL MNase (Worthington Biochem #LS004798) determined by the MNase titration was added. This digestion reaction was mixed at 37°C for 20 min on a thermomixer before being quenched with 4 mM EGTA (bioWORLD #40520008) and heat inactivated at 65°C for 10 min. Digested nuclei were washed twice with ice-cold Micro-C Buffer #2 (50 mM NaCl, 10 mM Tris-HCl pH = 7.5, 10 mM MgCl2, 100 µg/mL BSA (Sigma-Aldrich #B8667)).

**End repair and labeling**

As described in Goel *et al.* 2023, digested fragments were enzymatically blunted and biotinylated. First, digested chromatin was 5' phosphorylated in end-repair reactions (50 U T4 Polynucleotide Kinase (New England BioLabs #M0201), 50 mM NaCl, 10 mM Tris-HCl pH = 7.5, 10 mM MgCl2, 100 ug/mL BSA, 2 mM ATP (ThermoFisher #R1441), 5 mM DTT (Sigma-Aldrich #10197777001), in water) at 37°C for 15 min while mixing. Next, 50 U of DNA Polymerase I Klenow Fragment (New England BioLabs #M0210) was added to the reaction and incubated at 37°C for 15 min while mixing to create 5' fragment overhangs, and these overhangs were filled in by adding a mixture of dNTPs in end-labelling buffer (66 µM each of dTTP (Jena Bioscience #NU-1004), dGTP (Jena Bioscience #NU-1003), biotin-dATP (Jena Bioscience #NU-835-BIO14), and biotin-CTP (Jena Bioscience #NU-809-BIOX), 1X T4 DNA Ligase Buffer, 100 µg/mL BSA, in water) and incubating at room temperature for 45 min with interval mixing. This end-blunting reaction was quenched by 30 mM EDTA (Invitrogen #15575020) and heat inactivated at 65°C for 20 min. Finally, end-blunted and biotin-labeled nuclei were washed once with Micro-C Buffer #3 (50 mM Tris-HCl pH = 7.5, 10 mM MgCl2, 100 µg/mL BSA).

**Proximity ligation and removal of unligated biotin**

As described in Goel *et al.* 2023, proximity ligation was performed by incubating labeled chromatin in a ligation reaction (10,000 U T4 DNA Ligase (New England BioLabs #M0202), 1X T4 DNA Ligase Buffer, 100 µg/mL BSA, in 500 µL water) at room temperature for at least 2.5 hours or overnight with gentle mixing. To remove biotinylated dNTPs from all unligated fragment ends, samples were digested by 1,000 U of Exonuclease III (New England BioLabs #M0206) in reaction buffer (1X NEBuffer #1 in water) at 37°C for 15 min with interval mixing.

**DNA purification and size-selection**

As described in Goel *et al.* 2023, ligated DNA fragments were purified over a series of steps. DNA was first reverse crosslinked to remove proteins and RNA by adding 1% SDS (Sigma-Aldrich #L3771), 2 mg/mL Proteinase K (Viagen Biotech #501-PK), 250 mM NaCl, and 100 µg/mL RNaseA (ThermoFisher #EN0531) to the samples and incubating at 65°C overnight. Following crosslink reversal, the DNA solution was purified using the Zymo DNA Clean & Concentrator kit (Zymo Research #D4034) according to the kit manual.

Ligated DNA fragments were subsequently size-selected (~200-400 bp) by extraction from a 1% agarose gel (VWR #97062). Gel extracts were purified using the Zymo Gel Purification kit (Zymo Research #D4008), and samples were quantified by Qubit 1X dsDNA High Sensitivity Assay (Invitrogen #Q33231).

Ligated fragments were further purified by using Dynabeads MyOne Streptavidin T1 (Invitrogen #65601) to enrich for biotinylated fragments. DNA samples were bound to beads in a Binding and Wash Buffer (1 M NaCl, 5 mM Tris-HCl pH = 7.5, 500 µM EDTA, 0.1% Tween-20 (Sigma-Aldrich #P8074)) at room temperature for at least 30 minutes with mixing. After two washes with the Binding and Wash Buffer, the bead-bound samples were washed once with 10 mM Tris-HCl pH = 7.5 prior to library prep.

**Library preparation**

As described in Goel *et al.* 2023, Illumina library preparation was performed using the NEBNext Ultra II kit (New England BioLabs #E7645), with the addition of interval shaking (1 minute on, 3 minutes off) at 1000 rpm during incubations to mix the bead-bound samples. Sample washes were performed using Binding and Wash Buffer and 10 mM Tris-HCl pH = 7.5. A test library amplification determined the number of PCR cycles necessary to meet Capture input requirements (200-500 ng per sample) using

5% or less of the prepped library, with test PCR reactions run on an agarose gel and yields quantified using image quantification software Image Studio Lite (LI-COR Biosciences). The M-to-G1 RCMC replicates in this manuscript used 6-9 PCR cycles for final library amplification while the condensin degron RCMC replicates ranged from 7-17 cycles, with samples having RCMC inputs below 5M cells (e.g., all 0.5h and 8h replicates) requiring more cycles. Libraries were separately indexed by sample and replicate using sequencing indices from the NEB Multiplex Oligos for Illumina Primer Sets 1 and 2 (New England BioLabs #E7335 and #E7500), and amplification was done using the KAPA HiFi HotStart ReadyMix (Roche #07958927001). Following library amplification, amplified libraries were purified to remove adaptor dimers, primers, and contaminants using AmPure XP beads (Beckman Coulter #A63880). Purified libraries were quantified via Fragment Analyzer and qPCR at the MIT BioMicro Center to determine library concentrations for pooling prior to Capture.

**Capture probe design**

As described in Goel *et al.* 2023, target loci of interest were identified based on genomic features or enhancer-promoter relationships of interest (**Fig. S1b**). The *Klf1* locus, which we previously reported[4], was selected for its dense microcompartments. The *Dag1*, *Id1*, and *Cdt1* loci were selected as similarly gene-rich loci likely to exhibit a mixture of microcompartment, A/B compartment, and CTCF loop features while also containing genes relevant to cell cycle control. The *Myc* locus, heavily studied for MYC's role as a key transcription factor associated with disease and cell cycle control, was selected as a relatively gene-poor control with a well-characterized function and regulatory relationships. Using the UCSC Genome Browser and HiGlass visualization of existing G1E-ER4 Hi-C datasets, locus bounds were selected to include visible local structures and genomic features in roughly 1-2 Mb-sized regions. Once loci had been selected, 80-mer probes were designed to tile end-to-end without overlap across the Capture loci through Twist Bioscience. Probes with high predicted likelihoods of off-target pulldown (e.g., such as those in high-repeat regions) were masked and removed from the probe tiling, and probe coverage was double-checked to ensure the inclusion of key genomic features (e.g., all promoters and CTCF sites in the locus) before finalization. Probe panels were synthesized and purchased as Custom Target Enrichment Panels from Twist Bioscience.

**Capture of target loci**

As described in Goel *et al.* 2023, Capture was performed in accordance with Twist Bioscience's Standard Hybridization Target Enrichment Protocol. Sample libraries were pooled in a 1:1 molar ratio across conditions, after which they were dried and mixed with Hybridization Mix (Twist Bioscience #104178), Custom Panels (Twist Bioscience #101001), Universal Blockers (Twist Bioscience #100578), and Mouse Cot-1 DNA (Invitrogen #18440016). The library pool was hybridized to the biotinylated probe panel overnight, after which streptavidin beads (Twist Bioscience #100983) were used to pull down probes with hybridized ligated fragments and then washed (Twist Bioscience #104178) to remove unbound fragments. Another round of PCR amplified the target-enriched library using the Equinox Library Amplification Mix (Twist Bioscience #104178), with a test PCR included (as described above) to identify the number of amplification cycles necessary to meet sequencing requirements. With 2-4 µg of pooled input library for Capture, the RCMC samples generated in this manuscript needed 4-5 cycles of post-Capture PCR amplification. Following PCR amplification, the Captured library was purified (Twist Bioscience #100983) and then quantified via both Fragment Analyzer and qPCR at the MIT BioMicro Center in preparation for sequencing submission.

**Sequencing**

Following qPCR quantification, post-Capture libraries across replicates were pooled in a 1:1 molar ratio. Pooled libraries were paired-end sequenced using 2x50 cycle sequencing kits with Illumina NovaSeq S1 flow cells on a NovaSeq 6000 system (only Biological Replicate 1 of the M-to-G1 samples) or using 2x150 cycle sequencing kits on a NovaSeq X system (all biological replicates across all conditions) by the Broad Institute of MIT and Harvard's Walk-Up Sequencing services. Basecalls for NovaSeq output were performed using bcl2fastq v2.20.0.422.

### Data Analysis

##### Mapping and normalizing RCMC

RCMC paired-end reads generated by the Illumina NovaSeq sequencers were downloaded as .fastq files for each sample, pair mate, and flow cell lane. Read quality was verified using FastQC (v0.11.9). Paired end reads were aligned to the UCSC mm39 genome using bwa-mem2 (v2.2.1). Aligned paired end reads were then parsed with pairtools (v0.3.0) parse with --add-columns mapq --walks-policy mask --min-mapq 2. Parsed reads were filtered for PCR duplicates and unmapped/mutiple mapping reads with pairtools dedup with --max-mismatch 1. Remaining reads were indexed (pairix v0.3.7) and filtered (pairtools select) to retain only those reads where both read mates lie in a locus of interest (**Fig. S1b, 1c, 13a, 14b**). These filtered reads were subsequently converted to .cool format using cooler (v0.8.11) cload pairs, creating binned read counts across the genome for 50 bp bins. Finally, .cool files were converted to the .mcool format with cooler zoomify including the --balance option, compiling read counts for bins from 50 bp up to 10 Mb in size.

Contact matrices were balanced using iterative correction and eigendecomposition (ICE)[9] as previously described[4], which normalizes all rows and columns of a contact matrix sum to the same value. ICE balancing was performed on .mcool files containing data only within captured regions of interest (ROIs).

##### Visualizing RCMC

RCMC contact maps were visualized alongside genomic annotations and published ChIP-seq datasets using the HiGlass[10] browser (http://higlass.io/) and software (v0.8.0). Contact maps shown in figures were generated using cooltools[11] (v0.5.0) (https://cooltools.readthedocs.io/) (**Fig. 1a, 2a, 3a, 4b, 4e, Fig. S2d, 3, 4, 5, 6, 7a, 10, 11, 12**). Genomic tracks (i.e., ChIP-seq) and gene annotations for manuscript figures were generated using CoolBox[12] (v0.3.3). In generating our genomic tracks, we analyzed 23 public datasets (Supplementary Table 1) using processed bigWig files which were CrossMapped[13] (v0.6.1) (http://crossmap.sourceforge.net/) to the mm39 reference genome. Tracks were visualized using the Integrative Genomics Viewer (IGV)[14] (v2.10.3) to scale tracks by identifying local maxima and minimizing noise.

##### Replicate reproducibility analysis

The reproducibility of RCMC replicates (**Fig. S1e, 14a**) was evaluated using HiCRep[15] (v1.12.2) for contact maps at 5 kb resolution, with parameters lbr = 0 and ubr = 5000000. Reproducibility scores were calculated across regions for all replicates using the optimal h-value determined from a single replicate.

##### Comparing data across methods

Mapped sequencing reads were filtered using pairtools select to quantify read counts according to chosen evaluation criteria (**Fig. 1d, Fig. S1c, 2b, 2c, 13a, 14b**). Filtering was performed identically across the RCMC and Hi-C datasets on .pairs files containing mm39-mapped reads. RCMC .pairs files were generated as described above, while mm9-aligned .pairs files and loop calls containing all unique reads were downloaded for Hi-C (GSE129997) and CrossMapped to the mm39 genome.

Quantifications of read coverage across bins were calculated in Python using cooler to load unbalanced 250 bp resolution .cool files into memory as matrices. These matrices were then iterated through to determine the average number of interactions in each contact bin (**Fig. 1d**) and the fractions of bins containing at least one read at different contact distances (**Fig. S2b, 2c**).

Genome-wide equivalents for RCMC data were calculated by extrapolating the number of unique contacts mapped to a Capture locus to a region the size of the entire mouse genome. This approach assumes homogeneous read coverage throughout the genome; in reality, however, read coverage is unevenly distributed between regions depending on the specific region and which 3C method is used. Specifically, genome-wide Hi-C also had higher coverage at the *Ctd1* region than the genome-wide average. As such, compared to Hi-C at *Ctd1*, RCMC captured ~351-fold more unique contacts in the most deeply sequenced condition (late G1).

##### Contact decaying curve analysis

Contact decay curves were generated by plotting contact probability against genomic separation using cooltools[11] (**Fig. 1c, 4g, Fig. S2a**). Balanced and smoothed curves were generated using contact matrices across each chromosome binned to 150 bp resolution. RCMC curves were truncated at 1 Mb genomic separation due to noise at larger genomic separations (all loci lie between 1-2 Mb in size).

##### Downsampling

RCMC datasets were downsampled using pairtools sample to randomly select a subset of the mapped contact pairs. To downsample RCMC (**Fig. S2c, 2d**), a .pairs file containing all mapped reads across all replicates was downsampled using downsampling ratios corresponding to ten orders of two from 1/2 to 1/1024. Each downsampled .pairs output file was then filtered for reads with both mates within one of the five Captured loci, the unique reads were extracted, and a .mcool was generated for visualization and analysis.

##### Chromatin loop analysis

Chromatin loops were initially called on M-to-G1 RCMC data using Mustache[16] (v1.2.4) (https://github.com/ay-lab/mustache) at 0.25, 0.5, 1, 2, 5, and 10 kb data resolutions with sparsity thresholds of 0.7 and q-value thresholds of 0.1. Finer resolutions of loop calling identified more microcompartmental loops, but still missed many loops while also increasingly misidentifying stripes as loops, overlapping loops, and clustering calls at short genomic distances off of the diagonal.

Manual loop-calling was subsequently performed in an attempt to minimize these artifacts in microcompartment analysis. We defined loops as punctate foci of interaction (i.e., "dots"), visibly discernible as being enriched relative to their local background. We did not identify diffuse and overly faint interactions, homogeneously enriched stripes, and very short-range loops just off of the diagonal (i.e., under ~5 kb genomic separation) as loops. Loops were called on ICE-balanced M-to-G1 datasets to create a superset of interactions spanning prometaphase through late G1, with the ana/telophase and late G1 conditions serving as the primary datasets for manual annotation. Calling was done across data resolutions from 150 bp to 3.2 kb resolution using the HiGlass browser interface. Scale bar limits were dynamically modulated to minimize background and clearly distinguish focal enrichment. A total of 3350 focal dots (loops) spanning 363 loop anchors were manually annotated across the five loci (**Fig. 2d, 2e**).

Previously published M-to-G1 Hi-C loop calls (GSE129997) were downloaded and lifted over to mm39-aligned coordinates (**Fig. S7a**). Calls within the capture loci were merged across the five M-to-G1 conditions, with calls within 10 kb of one another merged into a single loop call with averaged coordinates to avoid redundancy. A total of 134 loops spanning 227 unique anchors were found across the five loci (**Fig. S7d, 7e**).

##### Loop anchor classification using "inclusive" and "exclusive" approaches

To classify loop anchors as promoter, enhancer, or CTCF and cohesin-bound (**Fig. 2d, Fig. S7d**), loop anchor locations were compared with the corresponding chromatin features as follows. Promoter regions were defined using all TSS locations in the mm39 UCSC RefGene annotation[17] ±2 kb. Enhancers or CTCF and cohesin-bound sites were defined based on overlap of H3K4me1 (GSM946535) and H3K27ac (GSE61349), or CTCF (GSE129997) and RAD21 (GSE129997), respectively. For all datasets, bigWig files were converted to bedgraph files using UCSC bigWigToBedGraph (v377)[18], followed by peak calling using MACS2 bdgpeakcall[19] (v2.2.7.1). For CTCF, called peaks were then overlapped with CTCF sites identified using FIMO (v5.4.1): first, fasta-get-markov was used to generate a background model using the mm39 genome assembly, then motifs were identified using –max-stored-scores 50000000 –thresh 1e-3. Finally, locations of motifs were overlapped with peaks identified in ChIP-seq data, and only the motif with the highest score for each peak was maintained. The peaks (H3K4me1, H3K27ac, RAD21) or identified sites (CTCF) were then overlapped using bedtools intersect (v2.30.0) to give enhancers or CTCF and cohesin-bound regions. Anchors of interactions ±1 kb were then overlapped with each of the three features to classify them as promoter, enhancer, or CTCF and cohesin-bound. Anchors overlapping none of these three features were classified as "Other".

In some case, anchor fit multiple categories, e.g., in cases where a *cis*-regulatory element (an enhancer or promoter) was also bound by CTCF and cohesin. To accommodate these cases, we use either an "inclusive" or "exclusive" classification depending on the analysis. For analyses where we used the "inclusive" classification, we took a hierarchical approach classifying all promoters as promoters even if they overlap other features, then classifying all enhancers as enhancers even if they overlap other features, leaving CTCF and cohesin anchors as those that are bound by CTCF and cohesin, but not overlapping promoters and enhancers. For analyses where we used the "exclusive" classification, any anchors that overlapped both E/P and CTCF/cohesin – and any loops formed by these anchors – were removed from consideration, limiting the analysis to "pure" anchors and loops representative of single rather than multiple organizational mechanisms.

We used the "inclusive" classification for all contact map loop overlays and the following figure panels: **Fig. 2b-f, Fig. S7b-e**. We used the "exclusive" classification for all loop strength calculations to avoid obfuscating the contribution of different organizational mechanisms towards the temporal dynamics of M-to-G1 loop formation. Specifically, the "exclusive" classification was used for the following figure panels: **Fig. 2g, 3b-e, 4c-d, Fig. S9, 13c, 13d**.

The number of interactions formed by each anchor and the lengths of the interactions they form (**Fig. 2b, 2c, 2f, Fig. S7b, 7c**) was determined and visualized in Python using the matplotlib package.

##### Heatmap and metaplot generation

Heatmaps and metaplots were generated for annotated loop anchors using deeptools[20] (v3.5.1) computeMatrix followed by plotHeatmap in a region ±1 kb around the center of each anchor for genomics data listed in Supplementary Table 1 (**Fig. S8**).

##### Calculation of background-subtracted loop strength in RCMC contact maps

The loop strength score is calculated as a sum of values of the iteratively-corrected Micro-C contact map minus local background within a 21x21 square window centered on the loop (**Fig. 2g, 3c-e, 4c, Fig. S9a, 13c**). At a resolution of 500 bp, this window encompasses interactions whose anchors are within ±5 kb from the loop. The local background matrix, which is also a 21x21 square window centered on the loop, is constructed such that each matrix element corresponding to a genomic separation *s* is the average of all matrix elements of genomic distance *s* within ±100 kb from the loop. The local background matrix is designed to represent the expected amount of distance-dependent random contacts in the absence of an active looping mechanism.

##### Pile-up and loop strength analysis

Pile-up visualizations and intensity quantifications of annotated looping interactions were performed using cooltools[11] to generate aggregate peak analyses (**Fig. 3b, 4d, Fig. S9b, 13d**) and individual loop strength plots (**Fig. 2g, 3c-e, 4c, Fig. S9a, 13c**). Plots for all loops of a given classification (e.g., E-P loops) were generated and analyzed individually or averaged for a 24 kb (**Fig. 3b, Fig. S9b**) or 20 kb (**Fig. 4d, Fig. S13d**) window centered on the loop at 250 bp resolution. Strength was calculated as observed signal divided by expected signal throughout the manuscript, with observed signal being calculated as the integral of ICE-balanced matrix values within 5 kb of the loop and expected signal being calculated as the local *P(s)* curve at the loop. The sole exception to the use of observed over expected strength calculations is **Fig. 3d**, in which simply the observed signal is considered to directly compare percent change between two conditions (ana/telophase and late G1) with notably different *P(s)* curves.

**Compartmentalization analysis**

Compartments were called by applying eigendecomposition to the 0h, 1h, and 4h condensin depletion RCMC contact matrices using cooltools[11] (**Fig. 4e, 4f**). RCMC data was first ICE-normalized to remove the distance-dependent effect of contact frequency. Eigendecomposition was then performed, with GC content serving as a correlate for orienting eigenvectors to indicate A- (gene-rich or active chromatin) or B- (gene-poor or inactive chromatin) compartments. Finally, eigenvectors were binarized and visualized as tracks and map overlays, as well as region-specific compartmentalization saddleplots, using cooltools[11]. Compartment calling and saddleplot generation was performed at 0.5, 1, 2, and 5 kb data resolutions, with all producing similar output (calls at 2 kb resolution are shown for clarity's sake).

### *Polymer Simulations and Analysis*

##### Polymer simulations

We performed polymer molecular dynamics simulations using custom-written code (https://github.com/mirnylab/microcompartments) that uses the polychrom library[21] (see https://github.com/open2c/polychrom/) which is a Python wrapper for the OpenMM molecular simulation toolkit[22,23]. These codes are freely and publicly available. Simulations are conducted by coupling 1D loop extrusion simulations to 3D polymer simulations. Genomic positions of loop extruders are used to determine which polymer sites are physically bridged at a given instant time. Between loop extrusion simulation steps, the chromatin polymer evolves under the constraints set by the loop extruders bridging polymer sites. Simulation parameters are given in **Tables 1** and **2**.

##### Modeling loop extrusion

We simulate $N$ loop extruders on a 1D lattice of length $L$=61600 genomic sites, where each site represents $\sigma$=0.5 kb of chromatin (also see **Table 1** for parameters). The mean distance between loop extruders is given by $d=L/N$; thus the linear density of extruders is $1/d$. In simulation sweeps, $d$=250 kb unless noted. Each loop extruder occupies (and in the 3D simulation, bridges) two genomic sites. Each loop extruder is loaded at two adjacent unoccupied sites. Subsequent to loading, each of the two components of the loop extruder may translocate away from the position at which it was loaded with probability $p$, which leads to a macroscopic loop growth velocity $v=2p\sigma/\tau_0$, where $\tau_0$ is the time between loop extrusion timesteps, taken to be 0.5 s. Following *in vitro* observations[24–28], $v$=1 kb/s unless noted. Extrusion a loop extruder component continues until: 1) the loop extruder is stochastically unloaded at rate $1/\tau_{\mathrm{res}}$, where $\tau_{\mathrm{res}}$ is the mean residence time, 2) the loop extruder component encounters another loop extruder component, or 3) if the loop extruder models cohesin, it encounters a properly oriented CTCF stall site and stalls with probability $q$=0.5. CTCF sites for the simulated *Dag1* region were annotated from intersection of CTCF and RAD21 ChIP-seq peaks in the mm39 genome assembly. CTCF sites were only included in mitosis-to-G1 transition simulations.

##### Modeling polymer dynamics

A chromosome was modeled as a chain of $L$=61600 monomeric subunits, each representing $\sigma$=0.5 kb of chromatin (also see **Table 1**). Consecutive subunits were connected by harmonic springs. Monomers also interacted by soft repulsive interactions ($E_{\mathrm{repel}}$=3 $k_BT$), modeling excluded volume. Monomers were assigned as one of three types (A, B, or C; **Fig. 5a**) and could interact through attractive interactions, with homotypic affinities of $\epsilon_A$=0, $\epsilon_B$=0.05 $k_BT$, and $\epsilon_C$=0.9 $k_BT$ and heterotypic affinities of 0 unless noted. As in previous studies[29,30], the interaction potential between monomers a distance $r$ apart was given by the smooth square-like potential:

$$U(r) = \begin{cases} E_{\mathrm{repel}}\left(1+\frac{1}{E_0}\left(\left(\frac{a_0}{a}r\right)^2-1\right)\left(\frac{a_0}{a}r\right)^{12}\right) \text{ for } r<a \\ -\epsilon_i\left(\frac{1}{E_0}\left(\left(\frac{r-(a+a^*)/2}{(a^*-a)/2}a_0\right)^2-1\right)\left(\frac{r-(a+a^*)/2}{(a^*-a)/2}a_0\right)^{12}+1\right) \text{ for } r>a \end{cases}$$

where $a$ is the monomer diameter (see below), $a^*=1.5a$, $a_0=\sqrt{6/7}$, $E_0=46656\,/\,823543$, and $i$ denotes the type of homotypic compartmental interaction, if applicable. Assignment of monomer types was based on analysis and annotation of the 1.925 Mb *Dag1* region, with C-type monomers (microcompartments) assigned based on non-CTCF ana/telophase loop calls. C-type regions were uniformly taken to be 1.5 kb long. In each chromosome, eight 3850 monomer *Dag1* regions were simulated, with 3850 neutral (A-type) monomer segments intercalating between *Dag* 1 regions. Simulation results were insensitive to whether isolated *Dag1* regions or *Dag1* repeats were simulated. In simulation sweeps, simulated chromosomes were confined to a spherical region at the prescribed (volumetric) density $\rho_{\mathrm{chr}}$=0.25 unless noted. Simulations were evolved using the OpenMM fixed timestep Langevin integrator with (polychrom variables) timestep=40 and collision_rate=0.01. Polymer simulations were evolved for 540 timesteps between loop extrusion update steps (see below for time calibration).

##### Equilibration, time calibration, and data collection

Equilibrium simulations (for parameter sweeps) were equilibrated by first evolving 1D loop extruder dynamics for $10^6$ extrusion time steps ($\gg\tau_{\mathrm{res}}$). 3D chromosomes with active loop extrusion were then equilibrated by simulating for >$30\cdot10^6$ polymer time steps (>$20\tau_{\mathrm{res}}$). Equilibration was assessed by inspection of contact frequency curves and contact maps. Data was then collected from 1500 time points over $27\cdot10^6$ polymer time steps. Data was collected for a minimum of 10 independently equilibrated simulations per condition. RCMC-like contact maps were generated using a contact radius of 4 monomers and bin size of 4 monomers (2 kb).

Polymer and loop extruder simulation times were calibrated to each other and experimental timescales by measuring the two-point mean-squared displacement and root-mean-squared radius of gyration of a 515 kb region in simulations without loop extrusion. These measurements were compared to experimental measurements of the *Fbn2* locus in mouse embryonic stem cells[31] under cohesin degradation conditions (4 h RAD21-mAID depletion); this resulted in a monomer size of $a$=25 nm. Together with the choice of loop growth speed of 1 kb/s, this procedure resulted in the selection of 540 polymer steps per loop extruder step, with each loop extruder time step equivalent to 0.5 s.

##### Modeling the mitosis-to-G1 transition

Simulations of the mitosis-to-G1 transition were performed by first equilibrating loop extruder dynamics for 36000 extrusion steps ($\approx5\tau_{\mathrm{res}}$ for simulated condensin II) and then performing full polymer simulations with extrusion in cylindrical confinement (4:1 aspect ratio) at density $\rho_{\mathrm{chr}}$=0.65 for 300 minutes (i.e., 36000 more extrusion steps with 540 polymer steps per extrusion step).

Ends of the chromosome were tethered to opposite ends of the cylinder. Simulations proceeded as described in the text, with data collection for prometaphase at times 0<$t$<10 min, ana/telophase at 20<$t$<30 min, early G1 at 55<$t$<65 min, mid G1 at 115<$t$<125 min, and late G1 at 235<$t$<245 min.

During prometaphase, for $t$<15 min, loop extruders were condensins I and II with respective residence times, $\tau_{res}^{CI}$=3 min and $\tau_{res}^{CII}$=1 h and mean separations $d^{CI}$=35 kb and $d^{CII}$=140 kb (unless otherwise noted). Over the time 15<$t$<17 min, the condensin I level transiently increased to $d^{CI}$=27 kb before gradually decreasing to zero over 17<$t$<20 min.

During ana/telophase, over the time 25<$t$<30 min, the cylinder height decreased by half, while the radius increased in a way that gradually decreased chromatin density to $\rho_{chr}$=0.45. Over 30<$t$<35 min, the strength of the cylindrical confining potential was gradually decreased to zero while a new spherical confining potential with radius corresponding to $\rho_{chr}$=0.25 was gradually strengthened from zero.

Additionally, at $t$=30 min, condensin II was removed, A/B compartment interactions were activated, and CTCFs were added to the simulation. At this time, we began loading cohesin loop extruders with $\tau_{res}^{cohesin}$=10 min to the simulation. Cohesin loading continued throughout the simulation of G1 such that the cohesin level increased linearly with time and peaked at the end of the simulation with mean separation $d^{cohesin}$=100 kb.

**Table 2** provides a list of parameters used in the model.

**Computation of loop/dot strengths**

Loop/dot strengths in simulations were computed similarly to as described in experiments. Each dot strength (**Fig. 5f**, right) was computed as the mean number of contacts within a 6 kb x 6 kb window centered on the microcompartment or CTCF site of interest. To obtain relative dot strengths (**Fig. 5e**, left), individual dot strengths were scaled by the average number of contacts at the corresponding genomic distance (results were minimally quantitatively altered by scaling by the local mean number of contacts instead).

#### Code and data availability

RCMC analysis code is available on GitHub at https://github.com/ahansenlab/RCMC_mitosis_analysis_code and polymer simulation code is also available on GitHub at https://github.com/mirnylab/microcompartments

Sequencing data is available at NCBI GEO under accession number GSE276657.

**Table 1. Default parameters in parameter sweeps.**

| Parameter | Value | Notes & References |
|---|---|---|
| Loop extruder residence time, $\tau_{res}$ | 500 s | Within observed range for condensin and cohesin[31–39]. |
| Loop extruder linear density, $1/d$ | 1 / 100 kb | Within observed ranges[31,40–42]. |
| Loop extruder speed, $v$ | 1 kb/s | Based on *in vitro* studies[24–28]. |
| Probability of cohesin stopping at CTCF, $q$ | 0 | CTCF ignored for parameter sweeps. |
| Monomer/lattice site genomic size | 0.5 kb | Selected for resolution of microcompartments. |
| Monomer physical size | 25 nm | Calculated as described in Methods and Gabriele *et al.* 2022[31]. |
| Chromosome length | 30.3 Mb | Practical decision. |
| Rouse scaling prefactor for 515 kb segment, $\Gamma$ | 0.0076 $\mu m^2/s^{1/2}$ | Gabriele *et al.* 2022[31]. |
| A-type compartment affinity | 0 $k_BT$ | Conventional selection (e.g. see Ref [29,43]) |
| B-type compartment affinity | 0.05 $k_BT$ | Selected by parameter sweep |
| C-type (micro)compartment affinity | 0.9 $k_BT$ | Selected by parameter sweep |
| Chromatin volume fraction, $\rho_{chr}$ | 0.25 | Ou *et al.* 2017[44]. |

**Table 2. Default parameters in mitosis-to-G1 transition simulations.**

| Parameter | Value | Notes & References |
|---|---|---|
| Condensin I residence time, $\tau_{res}^{CI}$ | 3 min | Based on FRAP[33,34]. |
| Condensin II residence time, $\tau_{res}^{CII}$ | 1 h | Models stably bound condensin II observed in experiments[33,34] and similar to previous simulations[45]. |
| Cohesin residence time, $\tau_{res}^{cohesin}$ | 10 min | Based on FRAP studies[32,36–39]. |
| Condensin I linear density in prometaphase, $1/d^{CI}$ | 1 / 35 kb | Within observed ranges[34,46–49]. |
| Peak condensin I linear density prior to ana/telophase, $1/d^{CI,*}$ | 1 / 27 kb | Estimated from Ref [33,34]. |
| Condensin II linear density, $d^{CII}$ | 1 / 140 kb | Within observed ranges[34,46–49]. |
| Cohesin linear density, $d^{cohesin}$ | 1 / 100 kb | Within observed ranges[31,40–42]. |
| Loop extruder speed, $v$ | 1 kb/s | Based on *in vitro* studies[24–28]. |
| Probability of cohesin stopping at CTCF, $q$ | 0.5 | Selected by parameter sweep |
| Monomer/lattice site genomic size | 0.5 kb | Selected for resolution of microcompartments |
| Monomer physical size | 25 nm | Calculated as described in Methods and Gabriele *et al.* 2022[31]. |
| Chromosome length | 30.3 Mb | Practical decision |
| Rouse scaling prefactor for 515 kb segment, $\Gamma$ | 0.0076 $\mu m^2/s^{1/2}$ | Gabriele *et al.* 2022[31]. |
| A-type compartment affinity | 0 $k_BT$ | Conventional selection (e.g. see Ref [29,43]) |
| B-type compartment affinity | 0.05 $k_BT$ | Selected by parameter sweep |
| C-type (micro)compartment affinity | 0.9 $k_BT$ | Selected by parameter sweep |
| Interphase chromatin density, $\rho_{chr}$ | 0.25 | Ou *et al.* 2017[44]. |
| Mitotic chromatin density, $\rho_{chr}$ | 0.65 | Between previous experimental[44,50,51] and simulated values[52]. |
| Prometaphase cylindrical confinement aspect ratio | 4 | Consistent with observations across literature |

**Supplementary Table 1. List of published datasets used in this paper.**

List of public datasets used in this paper for the cell line G1E-ER4.

| | Organism | Sample | Condition | Description | GEO / ENCODE# | Figures | Reference |
|---|---|---|---|---|---|---|---|
| 2 | | | prometaphase | | GSE129997 | EX5, EX6, EX8 | Zhang, H., Emerson, D.J., Gilgenast, T.G. et al. Chromatin structure dynamics during the mitosis-to-G1 phase transition. Nature 576, 158–162 (2019). https://doi.org/10.1038/s41586-019-1778-y |
| 3 | | | ana/telophase | | GSE129997 | EX5, EX6, EX8 | Zhang, H., Emerson, D.J., Gilgenast, T.G. et al. Chromatin structure dynamics during the mitosis-to-G1 phase transition. Nature 576, 158–162 (2019). https://doi.org/10.1038/s41586-019-1778-y |
| 4 | | ChIP, CTCF | early G1 | Architectural protein, loop extrusion factor | GSE129997 | EX5, EX6, EX8 | Zhang, H., Emerson, D.J., Gilgenast, T.G. et al. Chromatin structure dynamics during the mitosis-to-G1 phase transition. Nature 576, 158–162 (2019). https://doi.org/10.1038/s41586-019-1778-y |
| 5 | | | mid G1 | | GSE129997 | EX5, EX6, EX8 | Zhang, H., Emerson, D.J., Gilgenast, T.G. et al. Chromatin structure dynamics during the mitosis-to-G1 phase transition. Nature 576, 158–162 (2019). https://doi.org/10.1038/s41586-019-1778-y |
| 6 | | | late G1 | | GSE129997 | EX5, EX6, EX8 | Zhang, H., Emerson, D.J., Gilgenast, T.G. et al. Chromatin structure dynamics during the mitosis-to-G1 phase transition. Nature 576, 158–162 (2019). https://doi.org/10.1038/s41586-019-1778-y |
| 7 | | | asynchronous | | GSE129997 | 2, EX5, EX6, EX8 | Zhang, H., Emerson, D.J., Gilgenast, T.G. et al. Chromatin structure dynamics during the mitosis-to-G1 phase transition. Nature 576, 158–162 (2019). https://doi.org/10.1038/s41586-019-1778-y |
| 8 | | | prometaphase | | GSE129997 | 2, EX5, EX6, EX8 | Zhang, H., Emerson, D.J., Gilgenast, T.G. et al. Chromatin structure dynamics during the mitosis-to-G1 phase transition. Nature 576, 158–162 (2019). https://doi.org/10.1038/s41586-019-1778-y |
| 9 | | | ana/telophase | | GSE129997 | 2, EX5, EX6, EX8 | Zhang, H., Emerson, D.J., Gilgenast, T.G. et al. Chromatin structure dynamics during the mitosis-to-G1 phase transition. Nature 576, 158–162 (2019). https://doi.org/10.1038/s41586-019-1778-y |
| 10 | | ChIP, RAD21 | early G1 | Architectural protein, loop extrusion factor | GSE129997 | EX5, EX6, EX8 | Zhang, H., Emerson, D.J., Gilgenast, T.G. et al. Chromatin structure dynamics during the mitosis-to-G1 phase transition. Nature 576, 158–162 (2019). https://doi.org/10.1038/s41586-019-1778-y |
| 11 | | | mid G1 | | GSE129997 | EX5, EX6, EX8 | Zhang, H., Emerson, D.J., Gilgenast, T.G. et al. Chromatin structure dynamics during the mitosis-to-G1 phase transition. Nature 576, 158–162 (2019). https://doi.org/10.1038/s41586-019-1778-y |
| 12 | | | late G1 | | GSE129997 | 2, EX5, EX6, EX8 | Zhang, H., Emerson, D.J., Gilgenast, T.G. et al. Chromatin structure dynamics during the mitosis-to-G1 phase transition. Nature 576, 158–162 (2019). https://doi.org/10.1038/s41586-019-1778-y |
| 13 | G1E-ER4 | | asynchronous | | GSE129997 | EX5, EX6, EX8 | Zhang, H., Emerson, D.J., Gilgenast, T.G. et al. Chromatin structure dynamics during the mitosis-to-G1 phase transition. Nature 576, 158–162 (2019). https://doi.org/10.1038/s41586-019-1778-y |
| 14 | | | prometaphase | | GSE129997 | 2, EX5, EX6, EX8 | Zhang, H., Emerson, D.J., Gilgenast, T.G. et al. Chromatin structure dynamics during the mitosis-to-G1 phase transition. Nature 576, 158–162 (2019). https://doi.org/10.1038/s41586-019-1778-y |
| 15 | | | ana/telophase | | GSE129997 | 2, EX5, EX6, EX8 | Zhang, H., Emerson, D.J., Gilgenast, T.G. et al. Chromatin structure dynamics during the mitosis-to-G1 phase transition. Nature 576, 158–162 (2019). https://doi.org/10.1038/s41586-019-1778-y |
| 16 | | ChIP, RNA Pol II | early G1 | Transcription | GSE129997 | EX5, EX6, EX8 | Zhang, H., Emerson, D.J., Gilgenast, T.G. et al. Chromatin structure dynamics during the mitosis-to-G1 phase transition. Nature 576, 158–162 (2019). https://doi.org/10.1038/s41586-019-1778-y |
| 17 | | | mid G1 | | GSE129997 | EX5, EX6, EX8 | Zhang, H., Emerson, D.J., Gilgenast, T.G. et al. Chromatin structure dynamics during the mitosis-to-G1 phase transition. Nature 576, 158–162 (2019). https://doi.org/10.1038/s41586-019-1778-y |
| 18 | | | late G1 | | GSE129997 | 2, EX5, EX6, EX8 | Zhang, H., Emerson, D.J., Gilgenast, T.G. et al. Chromatin structure dynamics during the mitosis-to-G1 phase transition. Nature 576, 158–162 (2019). https://doi.org/10.1038/s41586-019-1778-y |
| 19 | | ChIP, H3K4me1 | asynchronous | Histone marker, enhancers | GSM946535 | 2, EX5, EX6, EX8 | Mouse ENCODE Consortium., Stamatoyannopoulos, J.A., Snyder, M. et al. An encyclopedia of mouse DNA elements (Mouse ENCODE). Genome Biol 13, 418 (2012). https://doi.org/10.1186/gb-2012-13-8-418 |
| 20 | | ChIP, H3K4me3 | asynchronous | Histone marker, active genes | ENCFF098DTA | 2, EX5, EX6, EX8 | Mouse ENCODE Consortium., Stamatoyannopoulos, J.A., Snyder, M. et al. An encyclopedia of mouse DNA elements (Mouse ENCODE). Genome Biol 13, 418 (2012). https://doi.org/10.1186/gb-2012-13-8-418 |
| 21 | | ChIP, H3K27ac | asynchronous | Histone marker, enhancers | GSE61349 | 2, EX5, EX6, EX8 | Dogan N, Wu W, Morrissey CS, Chen KB et al. Occupancy by key transcription factors is a more accurate predictor of enhancer activity than histone modifications or chromatin accessibility. Epigenetics Chromatin 2015; |
| 22 | | ChIP, H3K9me3 | asynchronous | Histone marker, repressive chromatin | ENCFF480HQQ | EX5, EX6, EX8 | Mouse ENCODE Consortium., Stamatoyannopoulos, J.A., Snyder, M. et al. An encyclopedia of mouse DNA elements (Mouse ENCODE). Genome Biol 13, 418 (2012). https://doi.org/10.1186/gb-2012-13-8-418 |
| 23 | | ChIP, H3K27me3 | asynchronous | Histone marker, repressive chromatin | ENCFF978WJA | EX5, EX6, EX8 | Mouse ENCODE Consortium., Stamatoyannopoulos, J.A., Snyder, M. et al. An encyclopedia of mouse DNA elements (Mouse ENCODE). Genome Biol 13, 418 (2012). https://doi.org/10.1186/gb-2012-13-8-418 |
| 24 | | ChIP, H3K36me3 | asynchronous | Histone marker, gene bodies | ENCFF452ACX | EX5, EX6, EX8 | Mouse ENCODE Consortium., Stamatoyannopoulos, J.A., Snyder, M. et al. An encyclopedia of mouse DNA elements (Mouse ENCODE). Genome Biol 13, 418 (2012). https://doi.org/10.1186/gb-2012-13-8-418 |

### SUPPLEMNTARY REFERENCES
